# SARS-CoV-2 N protein-induced Dicer, XPO5, SRSF3, and hnRNPA3 downregulation causes pneumonia

**DOI:** 10.1101/2023.10.03.560426

**Authors:** Yu-Wei Luo, Jiang-Peng Zhou, Hongyu Ji, Anqi Zheng, Xin Wang, Zhizheng Dai, Zhicheng Luo, Fang Cao, Xing-Yue Wang, Yunfang Bai, Di Chen, Yueming Chen, Qi Wang, Yaying Yang, Xinghai Zhang, Sandra Chiu, Ai-Long Huang, Kai-Fu Tang

## Abstract

Age is a major risk factor for coronavirus disease (COVID-19)-associated severe pneumonia and mortality; however, the underlying mechanism remains unclear. Herein, we investigated whether age-related deregulation of RNAi components and RNA splicing factors affects COVID-19 severity. Decreased expression of RNAi components (Dicer and XPO5) and splicing factors (SRSF3 and hnRNPA3) correlated with increased severity of COVID-19 and SARS-CoV-2 nucleocapsid (N) protein-induced pneumonia. N protein induced autophagic degradation of Dicer, XPO5, SRSF3, and hnRNPA3, repressing miRNA biogenesis and RNA splicing and inducing DNA damage, proteotoxic stress, and pneumonia. Dicer, XPO5, SRSF3, and hnRNPA3 were downregulated with age in mouse lung tissues. Older mice experienced more severe N protein-induced pneumonia than younger mice. However, treatment with a poly(ADP-ribose) polymerase inhibitor (PJ34) or aromatase inhibitor (anastrozole) relieved N protein-induced pneumonia by restoring Dicer, XPO5, SRSF3, and hnRNPA3 expression. These findings will aid in developing improved treatments for SARS-CoV-2-associated pneumonia.

---

Coronavirus disease (COVID-19), caused by severe acute respiratory syndrome coronavirus 2 (SARS-CoV-2), is highly heterogeneous, ranging from asymptomatic and mild to severe and fatal^1^. Although COVID-19 indiscriminately affects individuals of all age groups, severe pneumonia and mortality occur disproportionately in older individuals^2–5^. An age-dependent increase in disease severity was also observed in a mouse model of COVID-19^6^. However, the basis for age-related differences in the severity of COVID-19 is poorly understood.

DNA damage is a pivotal contributor to aging, affecting most aspects of the aging phenotype^7–9^. It is also a driver of COVID-19 pathogenesis. Specifically, DNA damage response facilitates SARS-CoV-2 infection by boosting transcription of the SARS-CoV-2 entry receptor angiotensin-converting enzyme 2 (ACE2)^10,11^. Additionally, it may promote SARS-CoV-2 replication, as inhibiting DNA damage response represses SARS-CoV-2 replication^12^. Moreover, DNA damage may increase COVID-19 severity by inducing cell senescence, which contributes to “cytokine storm” in patients with COVID-19^3,13^. DNA damage also causes cytosolic DNA accumulation, which stimulates proinflammatory cytokine production^7,14,15^. Finally, DNA damage induces natural killer group 2, member D (NKG2D) ligand expression. Receptor engagement by NKG2D ligands triggers cytolysis and the release of proinflammatory factors by natural killer and other immune cells^16,17^.

Loss of proteostasis, another hallmark of aging, causes unrestrained inflammasome activation, the accumulation of dysfunctional organelles, and cellular damage and is associated with several age-related morbidities^9,18,19^. Deregulation of proteostasis may contribute to COVID-19 pathogenesis, as SARS-CoV-2 infection disturbs cellular proteostasis, and proteostasis perturbation represses SARS-CoV-2 replication^20,21^.

Dicer, a key component of the RNA interference (RNAi) pathway, is downregulated in human and mouse adipose tissues and in *Caenorhabditis elegans* during aging^22^. Age-related deregulation of RNAi components might be associated with hallmarks of aging, as knockdown of RNAi components leads to DNA damage and probably affects proteostasis by increasing protein synthesis owing to the global downregulation of microRNAs (miRNAs)^23–25^. Additionally, the RNAi pathway may suppress SARS-CoV-2 replication via RNA degradation. Mechanistically, Dicer cleaves viral double-stranded RNAs (dsRNAs) to generate small interfering RNAs (siRNAs), which guide the sequence-specific degradation of viral RNAs^26,27^. As a countermeasure, viruses encode suppressors to antagonize RNAi by sequestering Dicer and/or dsRNAs^28,29^. However, whether age-related deregulation of RNAi components affects COVID-19 severity remains unknown.

Age-related upregulation or downregulation of splicing factors occurs in multiple tissues (blood, brain, muscle, skin, and liver) of different organisms (mice, rats, and humans) and is associated with genomic stability and proteostasis^30–34^. Knockdown of splicing factors leads to the accumulation of R-loop—a transcription-coupled DNA-RNA hybrid that can induce DNA damage^31–33^. Moreover, RNA splicing inhibition leads to the production of intron-containing truncated proteins, a subset of which have intrinsically disordered regions, form insoluble cellular condensates, and trigger a proteotoxic stress response^34^. Additionally, RNA splicing is essential for SARS-CoV-2 replication, as splicing inhibition represses viral replication^20^. However, the role of age-related deregulation of RNA splicing factors in SARS-CoV-2 pathogenesis has not been investigated.

We hypothesized that age-related deregulation of RNAi components and RNA splicing factors in lung tissues leads to DNA damage and proteotoxic stress, which may aggravate SARS-CoV-2-induced pneumonia. Here, we investigated whether aging-related deregulation of RNAi components and RNA splicing factors impacts COVID-19 severity. Additionally, we determined whether SARS-CoV-2 induces lung injury by regulating the expression of RNAi components and splicing factors. This study may shed new light on the pathogenesis of SARS-CoV-2-induced pneumonia, and may aid in developing improved prevention and treatment options for SARS-CoV-2-associated pneumonia.

## Results

### RNAi components and splicing factors are downregulated in patients with severe COVID-19

To investigate whether expression levels of RNAi components and splicing factors are associated with COVID-19 severity, we analyzed the published monocytic myeloid-derived suppressor cell (M-MDSC) RNA sequencing data from patients with severe or asymptomatic COVID-19^35^. mRNA levels of the RNAi components and splicing factors in M-MDSCs were lower in patients with severe COVID-19 than in those with asymptomatic COVID-19 (Extended Data Fig. 1a). By analyzing another set of published RNA sequencing data^36^, we found that levels of RNAi components and splicing factors were also lower in lung tissues from deceased COVID-19 patients than in those from individuals without COVID-19 (Extended Data Fig. 1b). Collectively, these results suggest that decreased expression of RNAi components and splicing factors is associated with increased COVID-19 severity.

### SARS-CoV-2 nucleocapsid (N) protein interacts with and induces autophagic degradation of Dicer, XPO5, SRSF3, and hnRNPA3

To investigate whether SARS-CoV-2 modulates the function of the RNAi pathway, a cellular reversal-of-silencing assay was performed^29^. SARS-CoV-2 suppressed RNAi (Extended Data Fig. 1c, d). Using an intron-containing luciferase reporter minigene^37^, we demonstrated that SARS-CoV-2 inhibited the splicing of this minigene (Extended Data Fig. 1e).

Next, to investigate whether SARS-CoV-2-encoded proteins are responsible for the inhibitory effect on RNAi and RNA splicing, a panel of plasmids expressing different proteins encoded by SARS-CoV-2 were co-transfected with reporter plasmids. We found that N protein repressed RNAi and RNA splicing (Extended Data Fig. 1f–h).

To understand the molecular mechanisms underlying the N protein-mediated regulation of RNAi and RNA splicing, putative protein interactors of N protein were retrieved from the Biological General Repository for Interaction Datasets (BioGRID)^38^. We observed that the RNAi components and splicing factors may interact with N protein (Extended Data Fig. 2a). Co-immunoprecipitation assays confirmed the interaction between N protein and Dicer, exportin 5 (XPO5), serine/arginine-rich splicing factor 3 (SRSF3), and heterogeneous nuclear ribonucleoprotein A3 (hnRNPA3) in human lung cancer (A549) and human normal lung epithelial (BEAS-2B) cell lines (Fig. 1a and Extended Data Fig. 2b). Consistent with reports that N protein is an RNA-binding protein^39–41^, RNA-immunoprecipitation assay revealed its association with pre-miRNA (Fig. 1b). Treatment with RNases disrupted the interaction between N protein and Dicer, XPO5, SRSF3, and hnRNPA3 (Fig. 1c), suggesting that these interactions are RNA-dependent.

**Figure 1.**
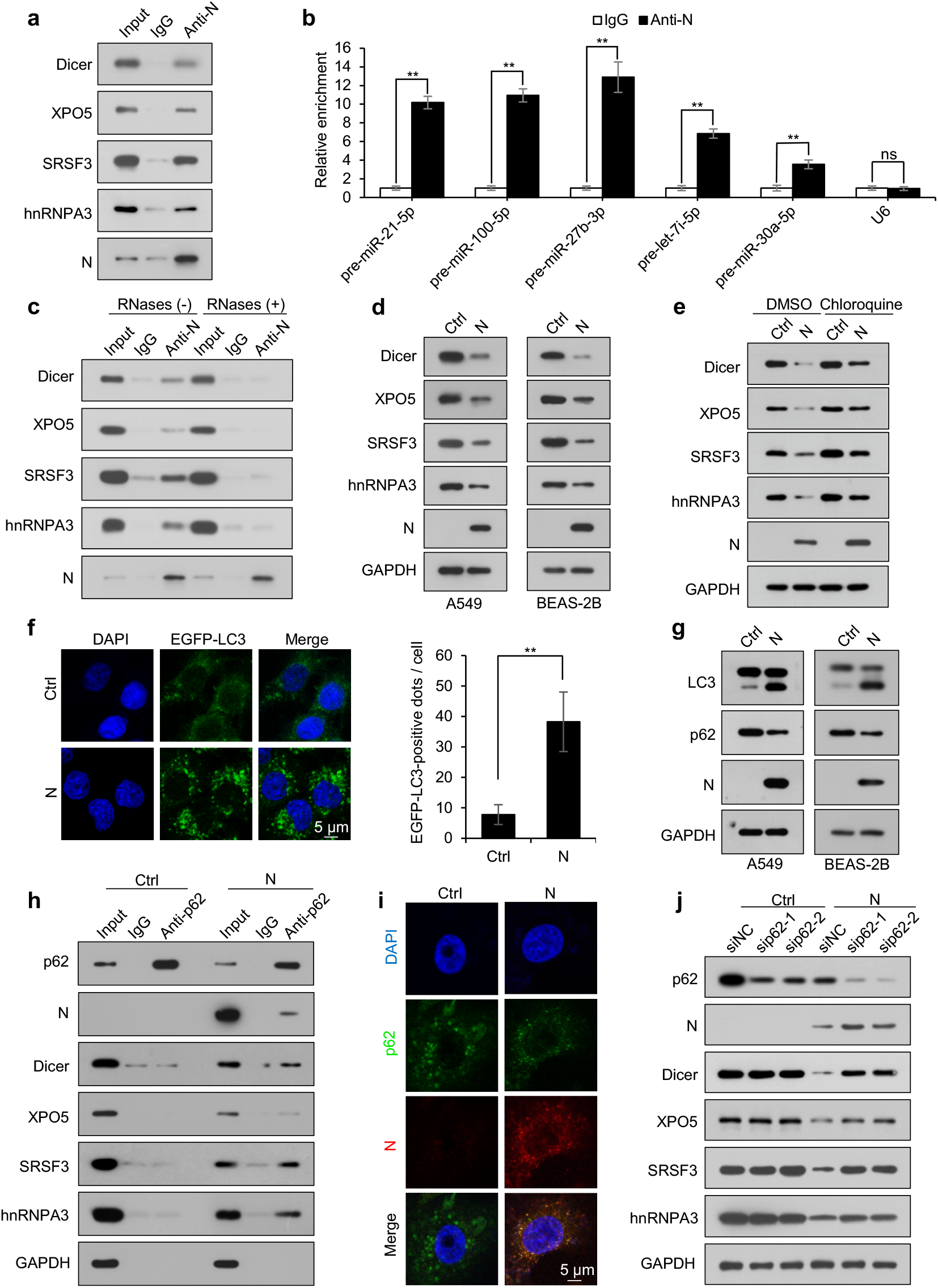
SARS-CoV-2 N protein interacts with and induces autophagic degradation of Dicer, XPO5, SRSF3, and hnRNPA3. See also Extended Data Fig. 1 and 2. **(a)** Lysates from A549 cells stably expressing N protein (A549-N) were subjected to immunoprecipitation using anti-N protein antibody and immunoblotting with the indicated antibodies. **(b)** Cell lysates of A549-N cells were subjected to RNA-immunoprecipitation (RIP) assay using anti-N protein antibody, followed by reverse transcription-quantitative polymerase chain reaction (RT-qPCR) detection of the indicated pre-miRNAs and *U6*. Data are expressed as mean ± standard deviation (SD) of three biological replicates. **p < 0.01; ns, not significant (p > 0.05; two-sided Student’s *t*-test). **(c)** Cell lysates of A549-N cells were subjected to immunoprecipitation using anti-N protein antibody in the presence or absence of RNases, followed by immunoblotting with the indicated antibodies. **(d)** Abundance of Dicer, XPO5, SRSF3, and hnRNPA3 proteins in A549 cells stably transfected with the control plasmid (A549-Ctrl), A549-N cells, BEAS-2B cells stably transfected with the control plasmid (BEAS-2B-Ctrl), and BEAS-2B cells stably expressing N protein (BEAS-2B-N). **(e)** Abundance of Dicer, XPO5, SRSF3, and hnRNPA3 proteins in A549-Ctrl or A549-N cells treated with or without chloroquine (20 µM). **(f**) The formation of EGFP-LC3 puncta in A549 cells stably expressing EGFP-LC3 transfected with N protein-expressing plasmid or control plasmid. Data are expressed as mean ± SD of three biological replicates, n ≥ 200 cells. **p < 0.01 (two-sided Student’s *t*-test). **(g)** LC3-II and p62 levels in A549-Ctrl and A549-N cells (left), or BEAS-2B-Ctrl and BEAS-2B-N cells (right). **(h)** Cell lysates of A549-Ctrl and A549-N cells were subjected to immunoprecipitation using an anti-p62 antibody and immunoblotting with the indicated antibodies. **(i**) Immunofluorescence analysis of A549-Ctrl or A549-N cells treated with chloroquine (20 µM) for 12 h, using anti-p62 (green) and anti-N protein (red) antibodies and DAPI (blue) for nuclear staining. **(j)** Immunoblotting analysis of the indicated proteins in A549-Ctrl and A549-N cells transfected with a negative control siRNA or p62-specific (sip62) siRNAs. Ctrl: control plasmid; N: N protein; anti-N: anti-N protein antibody; siNC: negative control siRNA.

Subsequently, we investigated whether N protein regulates Dicer, XPO5, SRSF3, and hnRNPA3 expression. Although N protein did not affect the mRNA levels of *Dicer, XPO5, SRSF3,* or *hnRNPA3* (Extended Data Fig. 2c), it decreased the abundance of their proteins in A549 and BEAS-2B cells (Fig. 1d). Treatment with an autophagy inhibitor, chloroquine, relieved the N protein-induced downregulation of Dicer, XPO5, SRSF3, and hnRNPA3 (Fig. 1e). However, treatment with a proteasome inhibitor, MG132, did not affect the N protein-induced downregulation of Dicer, XPO5, SRSF3, and hnRNPA3 proteins (Extended Data Fig. 2d), suggesting that the effect of N protein on the expression of Dicer, XPO5, SRSF3, and hnRNPA3 proteins is autophagy-dependent.

N protein was found to induce autophagy (Fig. 1f, g). A search of the BioGRID database indicated that N protein might interact with SQSTM1/p62, an autophagy receptor^42^. Co-immunoprecipitation and confocal microscopy analyses confirmed the interaction between N protein and p62 and their co-localization, respectively (Fig. 1h, i). Moreover, p62 was associated with Dicer, XPO5, SRSF3, and hnRNPA3 in N protein-expressing cells but not in cells without N protein expression (Fig. 1h). p62 knockdown partially relieved the N protein-induced downregulation of Dicer, XPO5, SRSF3, and hnRNPA3 proteins; however, it did not impact the Dicer, XPO5, SRSF3, or hnRNPA3 protein levels in cells not expressing N protein (Fig. 1j).

Furthermore, SARS-CoV-2 infection downregulated Dicer, XPO5, SRSF3, and hnRNPA3 at the protein but not the mRNA level (Extended Data Fig. 2e–g). Treatment with the autophagy inhibitor chloroquine relieved the SARS-CoV-2-induced downregulation of Dicer, XPO5, SRSF3, and hnRNPA3 proteins (Extended Data Fig. 2h).

Collectively, these findings indicate that N protein interacts with Dicer, XPO5, SRSF3, and hnRNPA3 proteins and induces their autophagic degradation in a p62-dependent manner.

### SARS-CoV-2 N protein induces DNA damage by downregulating Dicer, XPO5, SRSF3, and hnRNPA3 expression

As the RNAi machinery and splicing factors are essential for genomic stability^23,24,31–33^, we next investigated whether N protein induces DNA damage by downregulating Dicer, XPO5, SRSF3, and hnRNPA3 expression. Ectopic N protein expression induced DNA damage, as indicated by ATM, ATR, Chk1, Chk2, and H2AX phosphorylation, DNA break accumulation, and phosphorylated H2AX (γ-H2AX) foci formation (Fig. 2a–c and Extended Data Fig. 3a–c). Moreover, knockdown of Dicer, XPO5, SRSF3, or hnRNPA3 led to DNA damage (Fig. 2d, e and Extended Data Fig. 3d), whereas their overexpression partially alleviated the N protein-induced DNA damage (Fig. 2f, g and Extended Data Fig. 3e–h).

**Figure 2.**
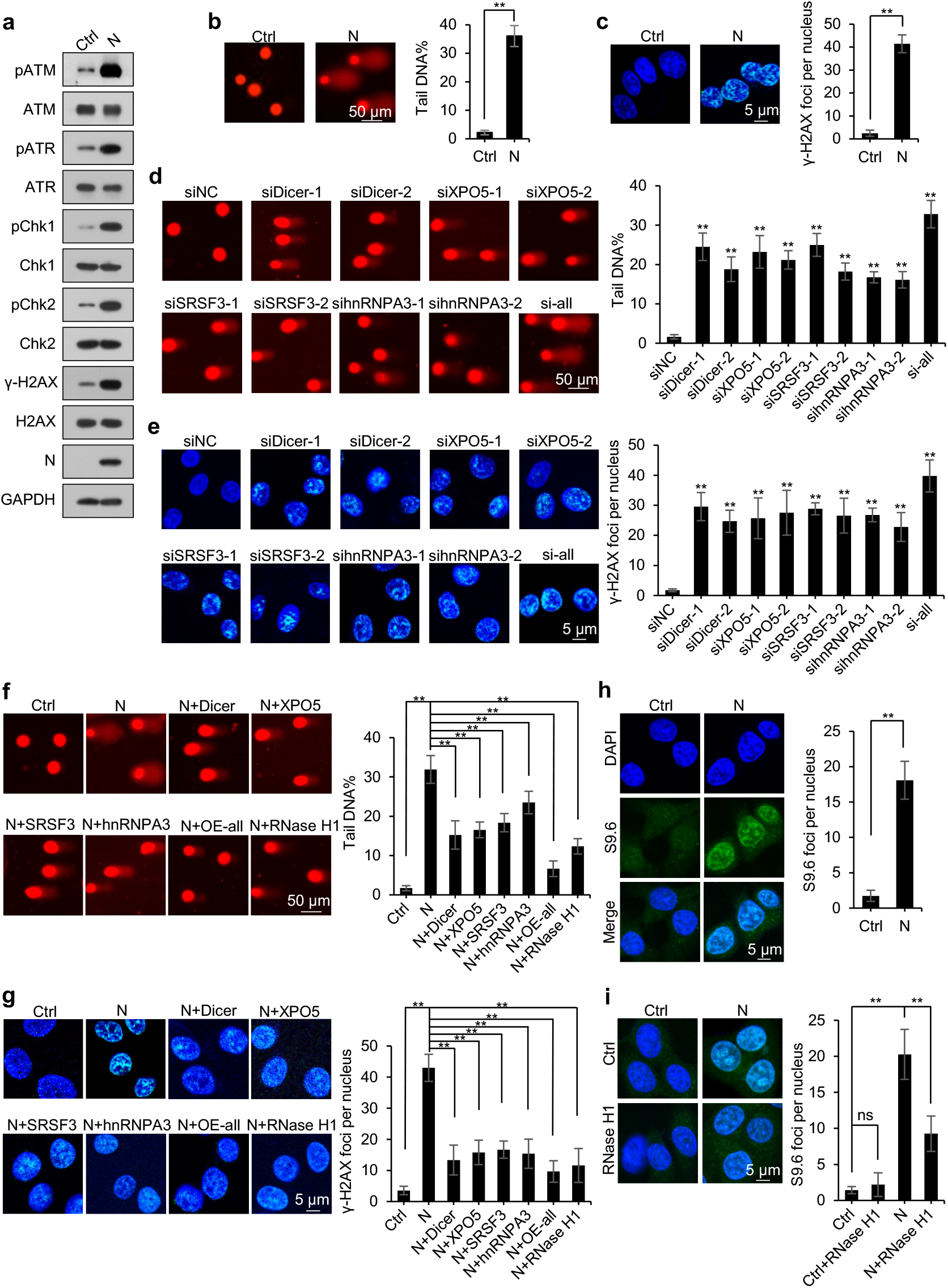
SARS-CoV-2 N protein induces DNA damage by downregulating Dicer, XPO5, SRSF3, and hnRNPA3 expression. See also Extended Data Fig. 3. **(a–c)** DNA damage in A549-Ctrl or A549-N cells was determined using immunoblotting analysis of the phosphorylation levels of ATM, ATR, Chk1, Chk2, and H2AX (a), comet assay (b), and immunofluorescence staining with anti-γ-H2AX antibody (c). **(d, e)** A549 cells were transfected with the indicated siRNAs and subjected to comet assay (d) and immunofluorescence staining with anti-γ-H2AX antibody (e). **(f, g)** A549-Ctrl and A549-N cells transfected with a control plasmid or plasmids overexpressing Dicer, XPO5, SRSF3, hnRNPA3, or RNase H1 were subjected to comet assay (f) and immunostaining with anti-γ-H2AX antibody (g). **(h)** Detection of R-loop in A549-Ctrl or A549-N cells using immunofluorescence staining with the S9.6 anti-DNA-RNA hybrid antibody. **(i)** Detection of R-loop in A549-Ctrl or A549-N cells transfected with a plasmid expressing RNase H1 or control plasmid using immunofluorescence staining with the S9.6 anti-DNA-RNA hybrid antibody. Data in b–i are expressed as mean ± SD of three biological replicates (n ≥ 200 cells). **p < 0.01; ns, not significant (p > 0.05) (two-sided Student’s *t*-test). Ctrl: control plasmid; N: N protein; siNC: negative control siRNA; si-all: cells transfected with siDicer, siXPO5, siSRSF3, and sihnRNPA3 together; OE-all: cells transfected with Dicer-, XPO5-, SRSF3-, and hnRNPA3-expressing plasmids together.

Defects in the splicing factors SRSF3 and hnRNPA3 increase R-loop formation, resulting in DNA damage^31–33^. We found that N protein induced R-loop accumulation (Fig. 2h); overexpression of RNase H1—an endoribonuclease that degrades the RNA portion of R-loop^43^—partially alleviated this effect and decreased DNA damage accumulation (Fig. 2f, g, i and Extended Data Fig. 3e–h). Overall, these findings indicate that N protein-induced downregulation of Dicer, XPO5, SRSF3, and hnRNPA3 expression results in DNA damage.

### SARS-CoV-2 N protein represses miRNA biogenesis

XPO5 and Dicer play pivotal roles in miRNA biogenesis. XPO5 transports the miRNA precursors (pre-miRNAs) from the nucleus to the cytoplasm, where Dicer processes them into mature miRNAs^23,25^. Small RNA deep sequencing revealed that N protein induced a global downregulation of miRNA expression (Fig. 3a and Extended Data Fig. 4a). Moreover, quantification of the top five miRNAs with the highest expression in A549 and BEAS-2B cells, accounting for approximately 80% of the entire cellular miRNome, revealed that they were all downregulated in N protein-expressing cells (Fig. 3b and Extended Data Fig. 4b).

**Figure 3.**
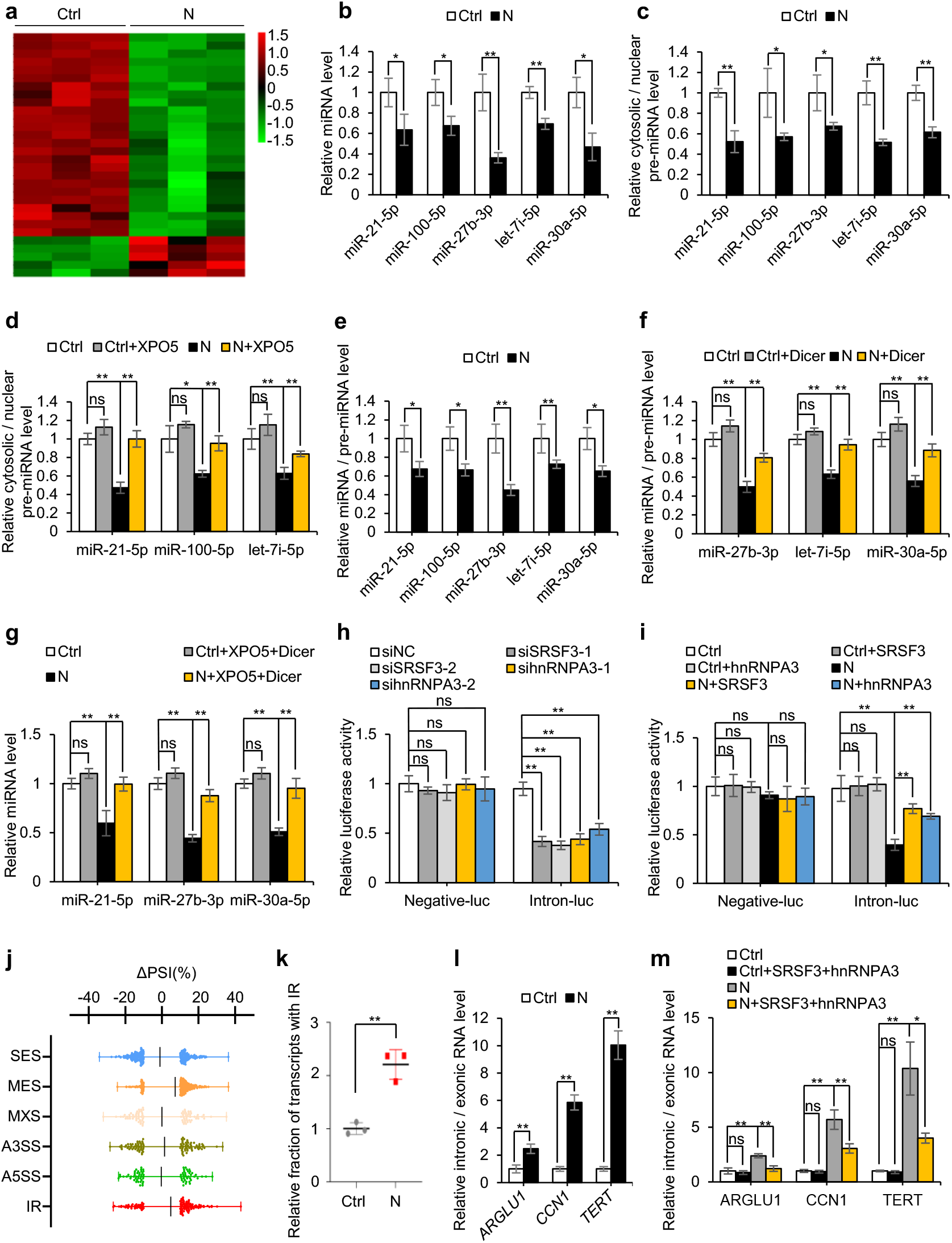
SARS-CoV-2 N protein represses miRNA biogenesis and splicing. See also Extended Data Fig. 4. **(a)** Heatmap of miRNA expression in A549-Ctrl and A549-N cells based on small RNA sequencing. **(b)** Quantification of five miRNAs in A549-Ctrl and A549-N cells. **(c)** Ratio of cytosolic pre-miRNA levels to nuclear pre-miRNA levels in A549-Ctrl and A549-N cells. **(d)** Ratio of cytosolic pre-miRNA levels to nuclear pre-miRNA levels in A549-Ctrl and A549-N cells transfected with a plasmid expressing XPO5 or control plasmid. **(e)** Ratio of mature miRNA levels to pre-miRNA levels in A549-Ctrl and A549-N cells. **(f)** Ratio of mature miRNA levels to pre-miRNA levels in A549-Ctrl and A549-N cells transfected with a plasmid expressing Dicer or control plasmid. **(g)** miRNA levels in A549-Ctrl or A549-N cells co-transfected with XPO5-expressing and Dicer-expressing plasmids. **(h)** Luciferase assay of HEK293T cells following co-transfection of splicing reporter with SRSF3-specific siRNA (siSRSF3), hnRNPA3-specific siRNA (sihnRNPA3), or control siRNA. **(i)** Luciferase assay of HEK293T-Ctrl or HEK293T-N cells following co-transfection of splicing reporter with SRSF3-expressing, hnRNPA3-expressing, or control plasmids. **(j)** Changes in percent spliced-in (PSI) values between A549-Ctrl and A549-N cells determined using full-length transcriptome sequencing data. ΔPSI = PSI(A549-N) − PSI(A549-Ctrl). **(k)** Relative fraction of transcripts with intron retention over total sequencing reads determined using full-length transcriptome sequencing data. **(l)** Relative intronic and exonic RNA levels of the indicated genes in A549-Ctrl and A549-N cells. **(m)** Relative intronic and exonic RNA levels of the indicated genes in A549-Ctrl and A549-N cells co-transfected with SRSF3-expressing and hnRNPA3-expressing plasmids. Data in b–i and k–m are expressed as mean ± SD of three biological replicates. **p < 0.01; *p < 0.05; ns, not significant (p > 0.05; two-sided Student’s *t*-test). Ctrl: control plasmid; N: N protein; siNC: negative control siRNA; SES: single exon skipping; MES: multiple exon skipping; MXE: mutually exclusive exons; A5SS or A3SS: alternative 5′ and 3′ splice sites, respectively; IR: intron retention.

Biochemical fractionation experiments revealed that N protein induced nuclear retention of pre-miRNAs, as indicated by an increase in pre-miRNA levels in the nucleus and their decrease in the cytoplasm (Fig. 3c and Extended Data Fig. 4c), which was relieved via XPO5 overexpression (Fig. 3d and Extended Data Fig. 4d). Moreover, N protein reduced the ratio between mature miRNA and pre-miRNA (Fig. 3e and Extended Data Fig. 4e), which was relieved by Dicer overexpression (Fig. 3f and Extended Data Fig. 4f). Dicer and XPO5 overexpression also rescued miRNA expression in N protein-expressing cells (Fig. 3g and Extended Data Fig. 4g).

Overall, these findings indicate that N protein blocks the transportation of pre-miRNAs from the nucleus to the cytoplasm by decreasing XPO5 expression and represses processing of pre-miRNAs into mature miRNAs by downregulating Dicer expression.

### SARS-CoV-2 N protein represses RNA splicing by decreasing SRSF3 and hnRNPA3 expression

Consistent with reports that SRSF3 and hnRNPA3 are splicing factors^32,33^, SRSF3 or hnRNPA3 knockdown inhibited the splicing of an intron-containing reporter minigene (Fig. 3h), whereas their overexpression partially relieved the inhibitory effect of N protein on RNA splicing (Fig. 3i). Transcriptome analysis revealed that N protein induced a global increase in intron retention (IR) (Fig. 3j, k). Additionally, reverse transcription-quantitative polymerase chain reaction (RT-qPCR) confirmed that N protein reduced the splicing efficiency of a panel of endogenous introns (Fig. 3l and Extended Data Fig. 4h). The inhibitory effect elicited by N protein on RNA splicing was partially alleviated by SRSF3 and hnRNPA3 overexpression (Fig. 3m). These results suggest that N protein represses RNA splicing by decreasing SRSF3 and hnRNPA3 expression.

### SARS-CoV-2 N protein induces proteotoxic stress and activates c-Jun N-terminal kinase

Splicing inhibition induces widespread IR and the subsequent translation of intron-retained mRNAs; a proportion of intron-derived polypeptides are condensation-prone and proteotoxic^34^. Metabolic labeling of nascent polypeptides with puromycin, followed by ultracentrifugation, revealed that N protein induced increased synthesis of insoluble proteins (Fig. 4a).

**Figure 4.**
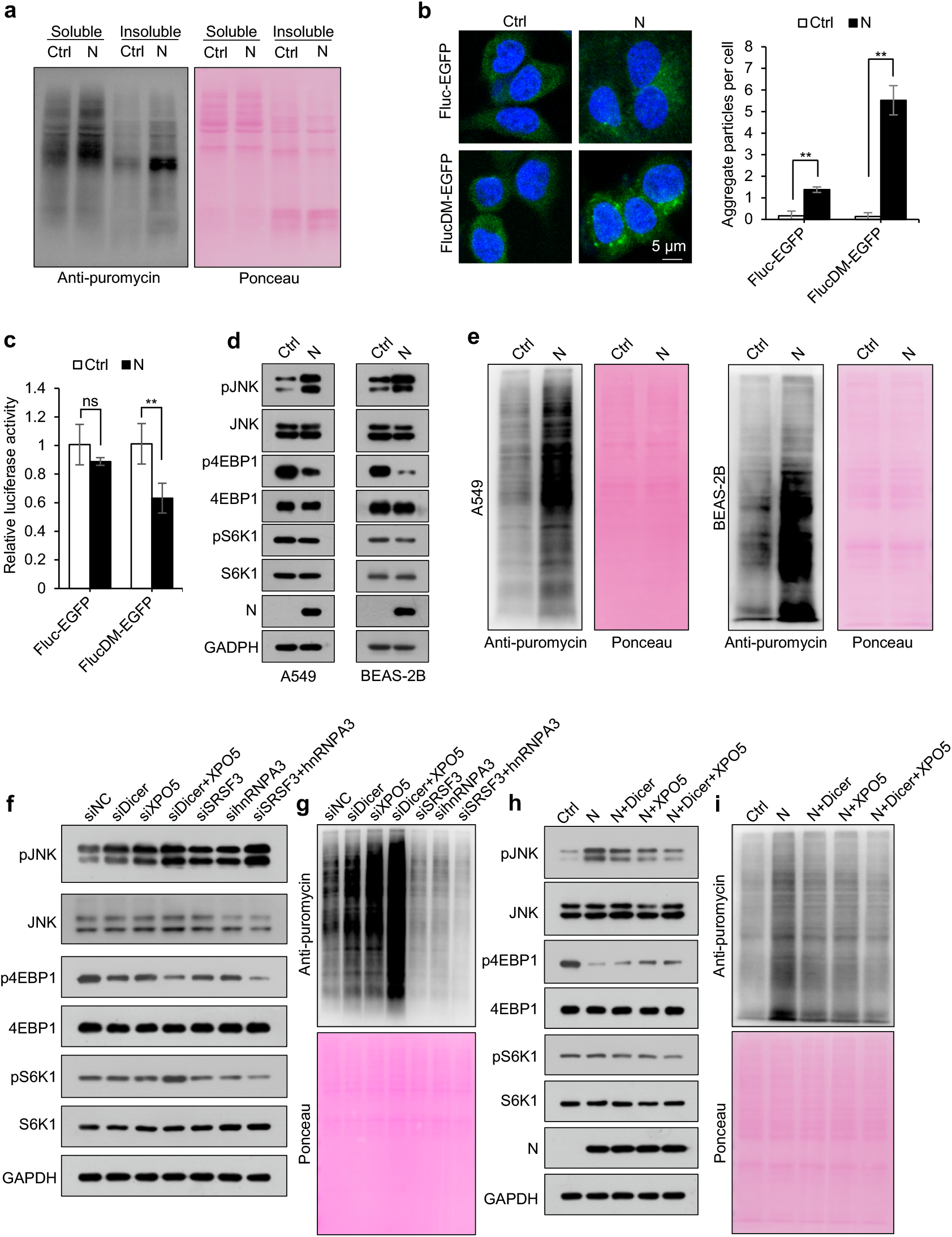
SARS-CoV-2 N protein induces proteotoxic stress and increases protein translation. **(a)** Immunoblotting of nascent polypeptides labeled with puromycin in the soluble and insoluble extracts of A549-Ctrl and A549-N cells. **(b, c)** A549-Ctrl or A549-N cells transfected with proteotoxic stress sensor reporters encoding Fluc-EGFP or FlucDM-EGFP and analyzed using confocal microscopy (b) or a luciferase assay (c). Data are expressed as mean ± SD of three biological replicates. **p < 0.01; ns, not significant (p > 0.05; two-sided Student’s *t*-test). **(d)** Immunoblotting of the indicated proteins in A549-Ctrl and A549-N (left) or BEAS-2B-Ctrl and BEAS-2B-N (right) cells. **(e)** Immunoblotting of nascent polypeptides labeled with puromycin in A549-Ctrl and A549-N cells (left) or BEAS-2B-Ctrl and BEAS-2B-N cells (right). **(f, g)** Immunoblotting of the indicated proteins (f) or nascent polypeptides labeled with puromycin (g) in A549 cells transfected with different siRNAs. **(h, i)** Immunoblotting of the indicated proteins (h) or nascent polypeptides labeled with puromycin (i) in A549-Ctrl, A549-N, and A549-N cells transfected with Dicer-expressing plasmid, XPO5-expressing plasmid, or both. Ponceau S staining image represents the loading control in a, e, g, and i. Ctrl: control plasmid; N: N protein; siNC: negative control siRNA.

To investigate whether N protein induces proteotoxic stress, we used a firefly luciferase (Fluc) or its conformationally unstable form (R188Q-R261Q double mutant [DM]), fused to enhanced green fluorescent protein (EGFP), as a reporter. Under proteotoxic stress, FlucDM-EGFP forms aggregates due to chaperone shortage^44^. Concordantly, N protein induced production of EGFP aggregates and reduced luciferase activity in cells harboring the FlucDM-EGFP reporter (Fig. 4b, c). Moreover, N protein activated c-Jun N-terminal kinase (JNK), a multifaceted stress-responsive kinase that can be activated by proteotoxic stress^34,45^, as evidenced by its increased phosphorylation (Fig. 4d). Overall, these findings suggest that N protein induces proteotoxic stress.

### SARS-CoV-2 N protein increases protein translation

Proteotoxic stress activates JNK, which induces disassembly of the mTORC1 complex, leading to dephosphorylation of eukaryotic translation initiation factor 4E binding protein 1 (4EBP1) and ribosomal protein S6 kinase beta-1 (S6K1) and repressing protein translation^34,45^. Although N protein led to 4EBP1 dephosphorylation, it increased protein translation without markedly impacting S6K1 phosphorylation (Fig. 4d, e).

Dicer or XPO5 knockdown led to JNK activation and 4EBP1 dephosphorylation while slightly increasing S6K1 phosphorylation and promoting protein synthesis. Simultaneous Dicer and XPO5 knockdown led to a more substantial increase in protein synthesis (Fig. 4f, g). Dicer and XPO5 overexpression partially prevented the N protein-induced increase in protein synthesis and JNK phosphorylation and caused a slight decrease in S6K1 phosphorylation and a slight increase in 4EBP1 phosphorylation in N protein-expressing cells (Fig. 4h, i). SRSF3 or hnRNPA3 knockdown activated JNK, dephosphorylated S6K1 and 4EBP1 and repressed protein synthesis (Fig. 4f, g). Overall, these findings indicate that N protein promotes protein translation by decreasing Dicer and XPO5 expression and inhibits protein synthesis by downregulating SRSF3 and hnRNPA3 expression.

### SARS-CoV-2 N protein induces pneumonia in mice

DNA damage and proteotoxic stress can induce cell death^7–9,18,19^. Consistently, N protein induced apoptosis in A549 cells (Extended Data Fig. 5a). Additionally, DNA damage induces accumulation of cytosolic DNA, which stimulates the expression of proinflammatory cytokines and NKG2D ligands^7,14–16^. Concordantly, N protein caused cytosolic DNA accumulation and promoted the expression of interferon beta (IFNβ), interleukin-6 (IL-6), and NKG2D ligands (Extended Data Fig. 5b, c).

We also investigated the *in vivo* effects of N protein and observed that its expression in mouse lung tissues led to DNA damage; cytosolic DNA accumulation; apoptosis; upregulation of IFNβ, IL-6, and NKG2D ligands; and pneumonia with obvious macrophage infiltration (Fig. 5a–d and Extended Data Fig. 5d, e). Given that N protein induced similar levels of lung injury in male and female mice (Extended Data Fig. 5f), we used only male mice for the following experiments. Dicer, XPO5, SRSF3, and hnRNPA3 knockdown in lung tissues led to lung injury and pneumonia, whereas their overexpression partially alleviated N protein-induced pneumonia (Fig. 5e, f and Extended Data Fig. 5g, h). Hence, N protein-induced downregulation of Dicer, XPO5, SRSF3, and hnRNPA3 protein expression is a mechanism underlying SARS-CoV-2-induced pneumonia.

**Figure 5.**
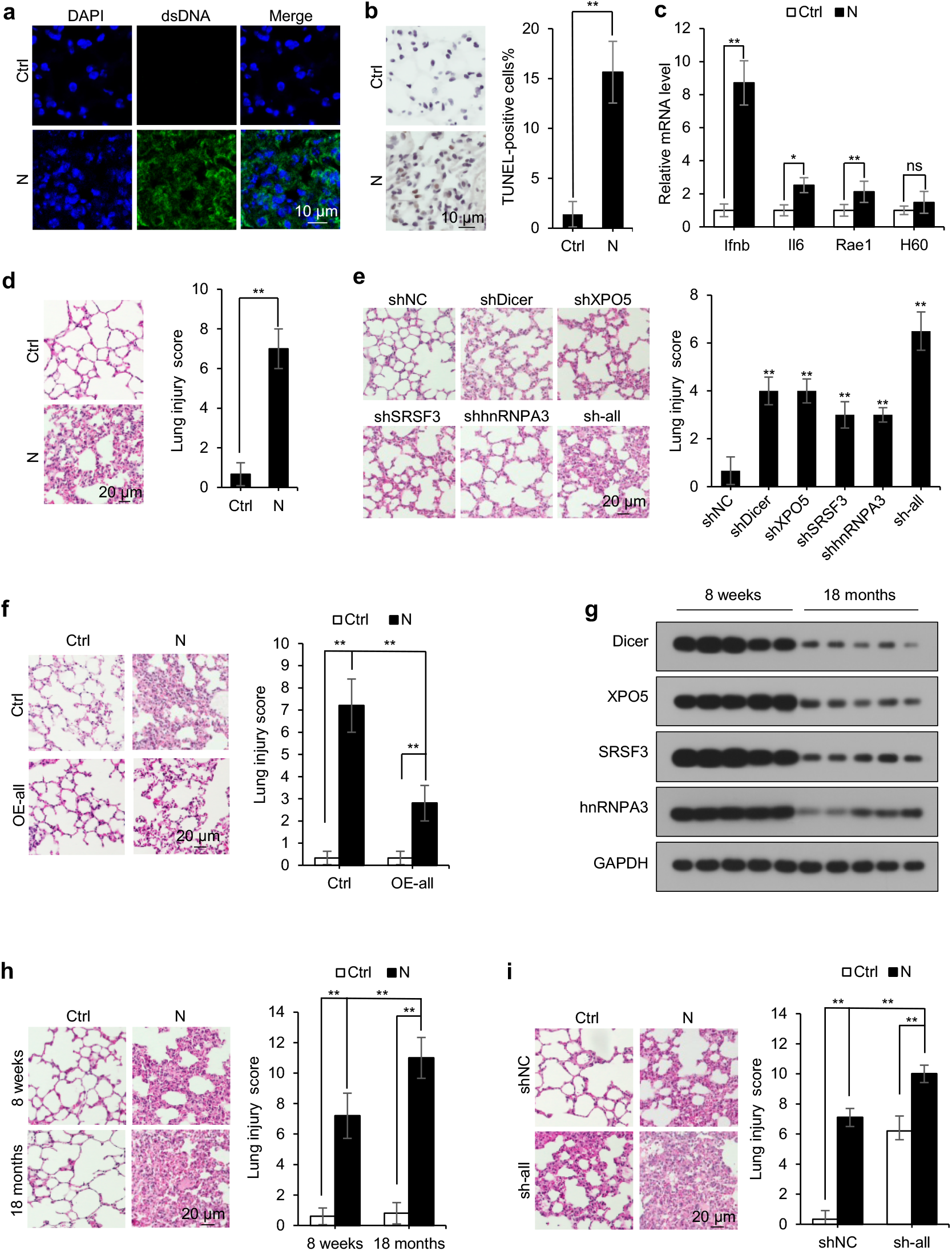
SARS-CoV-2 N protein induces pneumonia by downregulating Dicer, XPO5, SRSF3, and hnRNPA3 expression. See also Extended Data Fig. 5 and 6. **(a– d)** Eight-week-old male mice were intranasally instilled with control- or N protein-expressing plasmid, and lung tissues were subjected to immunofluorescence staining with anti-dsDNA antibody (a); terminal deoxynucleotidyl transferase dUTP nick end labeling (TUNEL) assay (b); RT-qPCR analysis of *Ifnb*, *Il6*, *Rae1*, and *H60* (c); and hematoxylin and eosin (HE) staining to assess lung injury (d). **(e)** Mice were intranasally instilled with plasmids expressing different shRNAs, and lung tissues were subjected to HE staining to determine the lung injury. **(f)** Mice were intranasally instilled with N protein-expressing plasmid or co-instilled with N protein-expressing plasmid and plasmids expressing Dicer, XPO5, SRSF3, and hnRNPA3, and lung tissues were subjected to HE staining to determine the lung injury. **(g)** Protein levels of Dicer, XPO5, SRSF3, and hnRNPA3 in the lung tissues of 8-week-old and 18-month-old mice. Mice (8-week-old and 18-month-old) were intranasally instilled with control or N protein-expressing plasmid and subjected to HE staining to determine the lung injury. Eight-week-old mice were intranasally instilled with N protein-expressing plasmid or co-instilled with N protein-expressing plasmid and shRNA-expressing plasmids; lung tissues were subjected to HE staining to determine the lung injury. Data in b–f, h and i are expressed as mean ± SD of five mice. **p < 0.01; *p < 0.05; ns, not significant (p > 0.05; two-sided Student’s *t*-test). Ctrl: control plasmid; N: N protein; shNC: negative control shRNA; sh-all: mice instilled with shDicer, shXPO5, shSRSF3, and shhnRNPA3 together; OE-all: mice instilled with Dicer-, XPO5-, SRSF3-, and hnRNPA3-expressing plasmids together.

### Age-related downregulation of Dicer, XPO5, SRSF3, and hnRNPA3 expression is associated with increased severity of N protein-induced pneumonia

Analyzing the published RNA sequencing data from lung tissues of 3-, 6-, 12-, and 24-month-old C57BL/6J mice revealed that mRNA levels of *Dicer, Xpo5, Srsf3,* and *Hnrnpa3* decreased with age (Extended Data Fig. 6a). The age-dependent downregulation of Dicer, XPO5, SRSF3, and hnRNPA3 in lung tissues was confirmed at the mRNA and protein levels in 8-week- and 18-month-old C57BL/6J mice (Fig. 5g and Extended Data Fig. 6b). Ectopic N protein expression in lung tissues led to DNA damage; apoptosis; cytosolic DNA accumulation; upregulation of IFNβ, IL-6, and NKG2D ligands; lung injury and pneumonia to a greater extent in older mice than in younger mice (Fig. 5h and Extended Data Fig. 6c–f). Furthermore, Dicer, XPO5, SRSF3, and hnRNPA3 knockdown exaggerated the N protein-induced DNA damage; apoptosis; cytosolic DNA accumulation; upregulation of IFNβ, IL-6, and NKG2D ligands; and pneumonia (Fig. 5i and Extended Data Fig. 6g–j). These results suggest that age-related downregulation of Dicer, XPO5, SRSF3, and hnRNPA3 is associated with increased severity of N protein-induced pneumonia.

### PJ34 alleviates N protein-induced pneumonia by preventing Dicer, XPO5, SRSF3, and hnRNPA3 downregulation

PJ34, a poly(ADP-ribose) polymerase (PARP) inhibitor, binds to N protein encoded by different coronaviruses and inhibits its RNA-binding activity^39–41^. PJ34 treatment inhibited the association between SARS-CoV-2 N protein and pre-miRNA and disrupted the interaction between N protein and Dicer, XPO5, SRSF3, and hnRNPA3 (Fig. 6a, b). Although PJ34 did not affect Dicer, XPO5, SRSF3, or hnRNPA3 expression in cells that did not express N protein, it increased their expression in N protein-expressing cells (Fig. 6c and Extended Data Fig. 7a).

**Figure 6.**
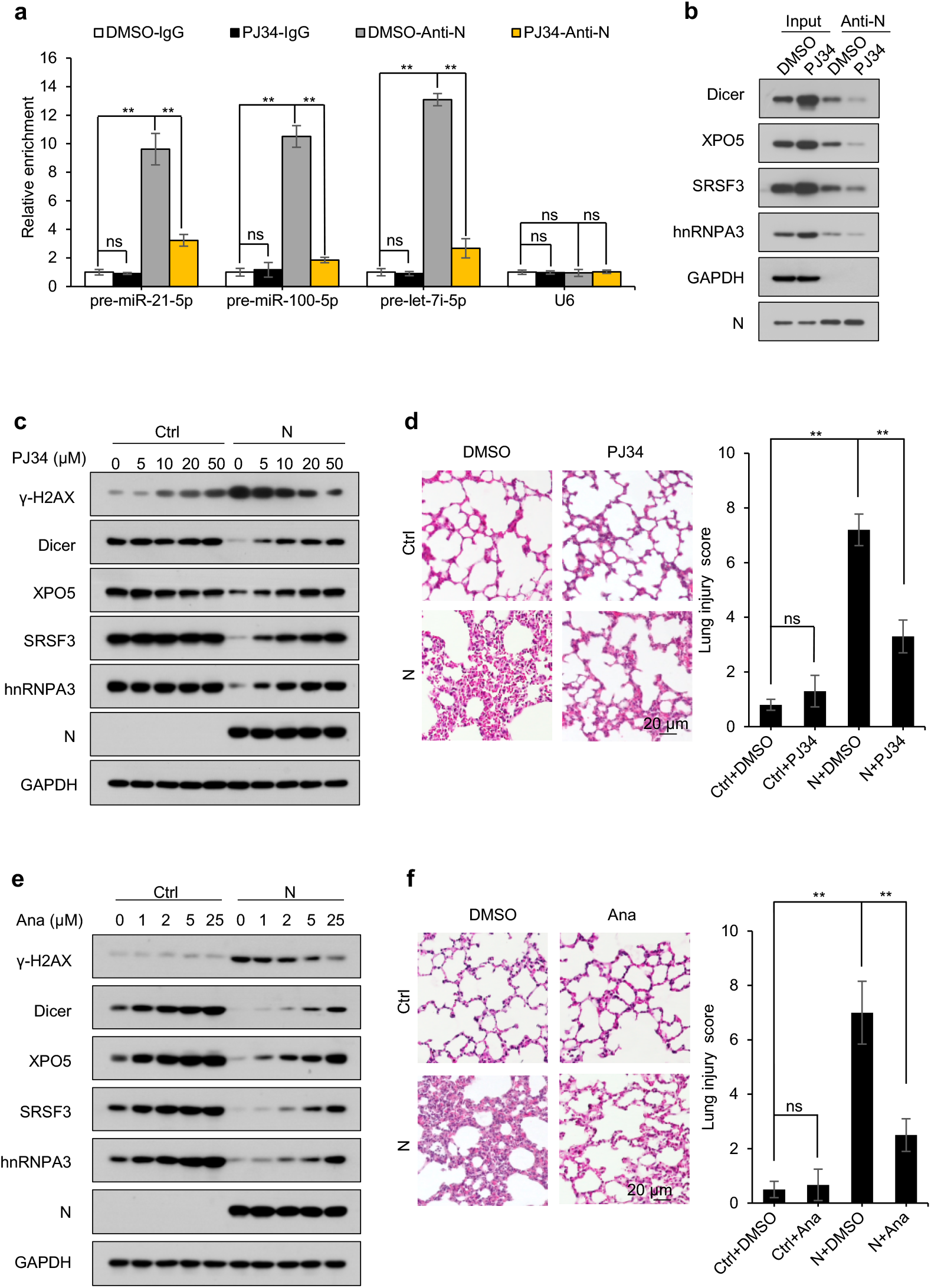
PJ34 and anastrozole relieve N protein-induced pneumonia. See also Extended Data Fig. 7. **(a, b)** Lysates from A549-N cells treated with or without PJ34 (50 µM) for 2 h were subjected to immunoprecipitation using anti-N protein antibody and then to RT-qPCR (a) and immunoblotting (b) to detect the pre-miRNAs and proteins that are associated with N protein. Data in a are expressed as mean ± SD of three biological replicates. **p < 0.01; ns, not significant (p > 0.05; two-sided Student’s *t*-test). **(c)** Immunoblotting of the indicated proteins in A549-Ctrl or A549-N cells after treatment with different concentrations of PJ34 for 2 h. **(d)** Mice were intranasally instilled with control plasmid or N protein-expressing plasmid and treated with or without PJ34 (10 mg/kg); lung tissues were subjected to HE staining to determine the lung injury. **(e)** Immunoblotting of the indicated proteins in A549-Ctrl or A549-N cells treated with different concentrations of anastrozole for 24 h. **(f)** Mice were intranasally instilled with control plasmid or N protein-expressing plasmid and treated with or without anastrozole (20 mg/kg); lung tissues were subjected to HE staining to determine the lung injury. Data in d and f are expressed as mean ± SD of five mice. **p < 0.01; ns, not significant (p > 0.05) (two-sided Student’s *t*-test). anti-N: anti-N protein antibody; Ctrl: control plasmid; N: N protein; Ana: anastrozole.

Although PJ34 induced minimal DNA damage in cells that did not express N protein, it alleviated the N protein-induced DNA damage (Extended Data Fig. 7a, b). Although PJ34 did not affect protein synthesis in cells not expressing N protein, it repressed the N protein-induced increase in protein synthesis and relieved the N protein-induced proteotoxic stress (Extended Data Fig. 7c, d). Moreover, PJ34 treatment ameliorated N protein-induced pneumonia (Fig. 6d). Collectively, these results indicate that PJ34 alleviates pneumonia by preventing the N protein-induced downregulation of Dicer, XPO5, SRSF3, and hnRNPA3.

### Anastrozole alleviates N protein-induced pneumonia by promoting Dicer, XPO5, SRSF3, and hnRNPA3 expression

Treatment with anastrozole, an aromatase inhibitor that enhances Dicer expression^46^, increased Dicer, XPO5, SRSF3, and hnRNPA3 expression in N protein-expressing cells and cells not expressing N protein (Fig. 6e and Extended Data Fig. 7e). Consistently, anastrozole treatment repressed the N protein-induced DNA damage accumulation, protein synthesis, and proteotoxic stress (Extended Data Fig. 7e–h) and ameliorated N protein-induced pneumonia (Fig. 6f). Collectively, these results reveal that anastrozole alleviates N protein-induced pneumonia by promoting Dicer, XPO5, SRSF3, and hnRNPA3 expression.

## Discussion

DNA damage and proteotoxic stress cause tissue injury by inducing cell death, inflammation, and tissue-destructive immune cell infiltration^7–9,14–16,18,19^. Herein, we demonstrated that SARS-CoV-2 N protein causes pneumonia by inducing DNA damage and proteotoxic stress. Mechanistically, N protein interacts with Dicer, XPO5, SRSF3, and hnRNPA3, leading to their autophagic degradation. Ectopic N protein expression induces DNA damage by downregulating Dicer, XPO5, SRSF3, and hnRNPA3 expression. Additionally, N protein disturbs proteostasis via two different mechanisms. First, it decreases SRSF3 and hnRPNA3 expression, repressing RNA splicing and inducing production of insoluble cellular condensates, triggering a proteotoxic stress response^34^. Second, it represses Dicer and XPO5 expression, eliciting a global increase in protein translation owing to a global reduction in miRNA levels^25^. Moreover, genetic or pharmacological rescue of Dicer, XPO5, SRSF3, and hnRNPA3 expression relieves the N protein-induced DNA damage, proteotoxic stress, and pneumonia.

Dicer, XPO5, SRSF3, and hnRNPA3 expression was downregulated in the lung tissues of older mice compared with that in younger mice, and N protein induced more severe lung injury and pneumonia in older mice. Dicer, XPO5, SRSF3, and hnRNPA3 knockdown exaggerated N protein-induced pneumonia. Therefore, age-associated downregulation of Dicer, XPO5, SRSF3, and hnRNPA3 expression in lung tissues may contribute to older individuals being more prone to developing severe pneumonia after SARS-CoV-2 infection.

The fundamental function of N protein is to package the viral genomic RNA into a ribonucleocapsid complex and promote viral RNA replication and transcription by recruiting host factors to the viral genome^47^. Our findings indicate that N protein causes pneumonia by inducing DNA damage and proteotoxic stress and may regulate SARS-CoV-2 replication via the following mechanisms. First, N protein suppresses RNAi by decreasing Dicer expression; therefore, it may promote SARS-CoV-2 replication by suppressing the antiviral RNAi^26,27^. Second, as DNA damage response inhibitors block SARS-CoV-2 replication^12^, N protein may promote SARS-CoV-2 replication by activating DNA damage response. Third, as splicing inhibition represses SARS-CoV-2 replication^20^, N protein may block SARS-CoV-2 replication by inhibiting RNA splicing. Fourth, although N protein inhibits IFN production via interaction with G3BP1^48,49^, we observed that it promotes IFNβ expression owing to cytosolic DNA accumulation induced by DNA damage. Therefore, N protein may promote or repress SARS-CoV-2 replication by downregulating or upregulating IFN expression, respectively. Finally, we observed that N protein can induce autophagy, which reportedly promotes or inhibits SARS-CoV-2 replication^50^. Therefore, N protein may regulate SARS-CoV-2 replication in an autophagy-dependent manner.

PJ34, a potent PARP inhibitor, elicits cytotoxicity against human cancer cells without impacting healthy cells^51^. It can also alleviate tissue damage and inflammation with different etiologies^52–54^. PJ34 binds to the coronavirus N protein, inhibiting its RNA-binding activity and hindering viral replication^39–41^. We found that PJ34 disrupted the interaction between SARS-CoV-2 N protein and Dicer, XPO5, SRSF3, and hnRNPA3 and relieved the N protein-induced downregulation of these proteins, alleviating the N protein-induced DNA damage and proteotoxic stress and ultimately mitigating N protein-induced pneumonia. Anastrozole, an aromatase inhibitor, is used to treat breast cancer and infertility^55–57^. We recently reported that anastrozole can treat colitis by promoting Dicer expression^46^. Here, we found that anastrozole alleviates N protein-induced pneumonia by promoting Dicer, XPO5, SRSF3, and hnRNPA3 expression. Therefore, it is worthwhile to conduct clinical trials to investigate whether PJ34 and anastrozole can be used to treat SARS-CoV-2-induced pneumonia.

Although the mRNA levels of *Dicer*, *XPO5*, *SRSF3*, and *hnRNPA3* were associated with COVID-19 severity, and their knockdown increased the severity of N protein-induced pneumonia in mice, whether their expression levels in pre-infected lung tissue are key determinants of COVID-19 severity remains unknown. Therefore, prospective studies in which human lung tissue samples are collected before SARS-CoV-2 infection are needed to determine whether pre-infection levels of Dicer, XPO5, SRSF3, and hnRNPA3 are associated with COVID-19 severity.

In summary, N protein leads to autophagic degradation of Dicer, XPO5, SRSF3, and hnRNPA3, inducing DNA damage and proteotoxic stress and eventually causing pneumonia. Treatment with PJ34 or anastrozole alleviates the N protein-induced DNA damage, proteotoxic stress, and pneumonia by rescuing Dicer, XPO5, SRSF3, and hnRNPA3 expression. The age-associated downregulation of Dicer, XPO5, SRSF3, and hnRNPA3 expression in lung tissues is one of the reasons why older individuals are more prone to developing severe pneumonia after SARS-CoV-2 infection than younger ones. Our findings can aid in developing improved treatments for SARS-CoV-2-associated pneumonia and imply that further studies should be conducted to investigate the toxicity and adverse events associated with N protein-based vaccines.

## Acknowledgments

We thank Dr. Yan Wu and Dr. Fangxu Li (Wuhan Institute of Virology of Chinese Academy of Sciences) for their assistance with SARS-CoV-2 virus preparation and infection. This work was supported by National Natural Science Foundation of China (grant numbers 81972648, 82172915, and 81773011), Chongqing Medical University Program for Youth Innovation in Future Medicine (grant number W0084), Science and Technology Innovation Project of Chongqing Medical University, and Chongqing Postdoctoral Science Foundation (grant number CSTB2023NSCQ-BHX0134).

## Author contributions

Conceptualization: K.F.T.; Methodology: K.F.T., Y.W.L.; Investigation: Y.W.L., J.P.Z., H.J., A.Z., X.W., Z.D., Z.L., F.C., X.Y.W., Y.B., D.C., Y,C., Q.W., Y. Y., X.Z., K.F.T.; Validation: Y.W.L., J.P.Z., H.J., A.Z., X.W., Z.D., Z.L., F.C., X.Y.W., Y.B., D.C., K.F.T.; Visualization: K.F.T., Y.W.L., J.P.Z., H.J.; Data curation: K.F.T., Y.W.L., J.P.Z., H.J.; Resources: X.Z., S.C., Funding acquisition: K.F.T., Y.W.L.; Project administration: K.F.T., Y.W.L.; Supervision: K.F.T., A.L.H.; Writing—original draft: K.F.T., Y.W.L.; Writing—review & editing: K.F.T.

## Competing interests

The authors declare no competing interests.

## Additional information

**Correspondence and requests for materials** should be addressed to Kai-Fu Tang.

## Online Methods

### Cell lines

The human lung adenocarcinoma epithelial cell line A549 (male, CCL-185), human bronchial epithelial cell line BEAS-2B (male, CRL-9609), and human embryonic kidney cell line HEK293T (female, CRL-3216) were obtained from the American Type Culture Collection (Manassas, VA, USA). A549 and BEAS-2B cells were cultured in Roswell Park Memorial Institute (RPMI)-1640 medium (Hyclone, Logan, UT, USA) supplemented with 10% (v/v) fetal bovine serum (FBS; Hyclone). HEK293T cells were cultured in Dulbecco’s modified Eagle’s medium (DMEM; Hyclone) supplemented with 10% (v/v) FBS. All cell lines were cultured at 37 °C in a 5% CO_2_ humidified incubator. All cell lines were routinely tested for *Mycoplasma* contamination using the *Mycoplasma* Detection Kit (Southern Biotech, Birmingham, AL, USA) and authenticated based on polymorphic short-tandem repeat loci. For the reporter gene experiments, we used HEK293T cells owing to their high transfection efficiency. For other experiments, A549 and BEAS-2B cells were used.

### Viruses

SARS-CoV-2 WIV04 (GISAID accession No. The EPI_ISL_402124) strain was originally isolated from COVID-19 patients and stored at the Wuhan Institute of Virology, Chinese Academy of Sciences^58^. All studies on live SARS-CoV-2 were performed under biosafety level-3 (BSL3) conditions at the Wuhan Institute of Virology, Chinese Academy of Science.

### Animals

Unless otherwise specified, 8-week-old or 18-month-old C57BL/6J male mice were used in this study. Five mice were included per group in each experiment. All mice were housed under pathogen-free conditions. The animal experiments were performed according to protocols approved by the Institutional Animal Care and Use Committee of the Wenzhou Medical University.

### Cell treatment

To block autophagy, the cells were treated with chloroquine (20 µM, MedChemExpress, Monmouth Junction, NJ, USA) for 12 h. To inhibit proteasome activity, the cells were treated with the proteasome inhibitor MG132 (20 µM, MedChemExpress) for 12 h. To investigate the effect of PJ34 on the interaction between N protein and Dicer, XPO5, SRSF3, or hnRNPA3, as well as their expression, cells were treated with PJ34 (MedChemExpress) at indicated concentrations for 2 h. To investigate the effect of anastrozole on the expression of Dicer, XPO5, SRSF3, and hnRNPA3, cells were treated with anastrozole (Aladdin, Shanghai, China) at the indicated concentrations for 24 h. Dimethyl sulfoxide (DMSO) was used as the solvent for chloroquine, MG132, PJ34, and anastrozole and served as the vehicle control.

### Mouse treatments

To ectopically express the genes of interest in mouse lung tissues, C57BL/6J mice were intranasally instilled with plasmids (20 μg per mouse) mixed with an *in vivo* transfection reagent (Entranster^TM^-in vivo, Engreen, Beijing, China) in a total volume of 40 μL. To knock down the genes of interest in mouse lung tissues, mice were intranasally instilled with shRNA plasmids (40 μg per mouse) mixed with the *in vivo* transfection reagent (Entranster^TM^-in vivo) in a total volume of 40 μL. The plasmids are listed in Extended Data Table 1. To investigate the role of PJ34 or anastrozole in N protein-induced pneumonia, 8-week-old male C57BL/6J mice were intraperitoneally injected with PJ34 (10 mg/kg) or anastrozole (20 mg/kg) at 12, 24, and 48 h after plasmid instillation. The mice were anesthetized using isoflurane (RWD Life Science, Shenzhen, China) and euthanized by cervical dislocation at 72 h after plasmid instillation, and the lung tissues were subjected to further analysis.

To investigate the effects of N protein on mouse lung tissues, 8-week-old male mice were randomly allocated to two groups (n = 5/group): group 1 was instilled with a control plasmid and group 2 was instilled with an N protein-expressing plasmid.

To investigate whether N protein exhibited different effects in male and female mice, 8-week-old male and female mice were allocated to four groups (n = 5/group): group 1, male mice were instilled with control plasmid; group 2, female mice were instilled with control plasmid; group 3, male mice were instilled with N protein-expressing plasmid; group 4, female mice were instilled with N protein-expressing plasmid.

To investigate whether N protein impacted old and young mice differently, 8-week-old and 18-month-old male mice were allocated to four groups (n = 5/group): group 1, 8-week-old mice were instilled with control plasmid; group 2, 18-month-old mice were instilled with control plasmid; group 3, 8-week-old mice were instilled with N protein-expressing plasmid; and group 4, 18-month-old mice were instilled with N protein-expressing plasmid.

To investigate whether Dicer, XPO5, SRSF3, or hnRNPA3 knockdown induced lung injury, 8-week-old male mice were allocated to six groups (n = 5/group) and instilled with control shRNA plasmid (shNC), Dicer shRNA plasmid (shDicer), XPO5 shRNA plasmid (shXPO5), SRSF3 shRNA plasmid (shSRSF3), hnRNPA3 shRNA plasmid (shhnRNPA3), or instilled with all four shRNA plasmids.

To investigate whether knockdown of Dicer, XPO5, SRSF3, or hnRNPA3 exaggerated N protein-induced pneumonia, 8-week-old male mice were allocated to four groups (n = 5/group): group 1, instilled with control plasmid and shNC; group 2, instilled with N protein-expressing plasmid and shNC; group 3, instilled with control plasmid, shDicer, shXPO5, shSRSF3, and shhnRNPA3; and group 4, instilled with N protein-expressing plasmid, shDicer, shXPO5, shSRSF3, and shhnRNPA3.

To investigate whether the overexpression of Dicer, XPO5, SRSF3, and hnRNPA3 alleviated N protein-induced pneumonia, 8-week-old male mice were allocated to four groups (n = 5/group): group 1, instilled with control plasmid; group 2, instilled with N protein-expressing plasmid; group 3, instilled with Dicer-, XPO5-, SRSF3-, and hnRNPA3-expressing plasmids; and group 4, instilled with N protein-, Dicer-, XPO5-, SRSF3-, and hnRNPA3-expressing plasmids.

To investigate whether PJ34 alleviated N protein-induced pneumonia, 8-week-old male mice were allocated to four groups (n = 5/group): group 1, instilled with control plasmid and intraperitoneally injected with DMSO; group 2, instilled with control plasmid and intraperitoneally injected with PJ34; group 3, instilled with N protein-expressing plasmid and intraperitoneally injected with DMSO; and group 4, instilled with N protein-expressing plasmid and intraperitoneally injected with PJ34.

To investigate whether anastrozole alleviated N protein-induced pneumonia, 8-week-old male mice were allocated to four groups (n = 5/group): group 1, instilled with control plasmid and intraperitoneally injected with DMSO; group 2, instilled with control plasmid and intraperitoneally injected with anastrozole; group 3, instilled with N protein-expressing plasmid and intraperitoneally injected with DMSO; and group 4, instilled with N protein-expressing plasmid and intraperitoneally injected with anastrozole.

### Histological analysis of acute lung injury

To assay lung injury, the lung tissues were fixed in 4% formaldehyde and dehydrated, embedded in paraffin, and sectioned at 4 μm thickness. The lung sections were stained with hematoxylin and eosin (HE) according to standard protocols, dehydrated in a graded series of ethanol, cleared in xylene, cover-slipped, and photographed using a light microscope (Axioscope 5; Carl Zeiss, Jena, Germany).

Acute lung injury (ALI) scoring was conducted by two pathologists blinded to the treatment protocol using the following parameters as previously described^59,60^: (1) septal mononuclear cell/lymphocyte/neutrophil/macrophage infiltration; (2) septal hemorrhage and congestion and alveolar hemorrhage; (3) alveolar edema and proteinaceous debris filling the airspace; (4) alveolar septal thickening. The severity of each category was graded from 0 (minimal) to 3 (maximal), and the total score was calculated by adding the scores for each category.

### Immunohistochemistry and immunofluorescent staining

Heat-induced antigen retrieval on lung tissue sections was performed in antigen retrieval solution. Endogenous peroxidase activity was blocked with 3% H_2_O_2_ (ZSGB-BIO, Beijing, China) for 10 min, followed by incubation with blocking solution (ZSGB-BIO) for 15 min to minimize non-specific staining. The slides were incubated with anti-CD3 (17617-1-AP; Proteintech, Chicago, IL, USA, 1:200), anti-CD22 (21894-1-AP; Proteintech, 1:2000), anti-CD68 (28058-1-AP; Proteintech, 1:1000), anti-Ly6G (ab238132; Abcam, Boston, MA, USA, 1:100), and anti-dsDNA (MAB1293; Sigma-Aldrich, St. Louis, MO, USA, 1:500) at 4 °C overnight.

For immunohistochemical staining, the slides were washed three times with phosphate-buffered saline (PBS) to remove unbound primary antibodies, then incubated with Elivision^TM^ super HRP (MXB Biotechnologies, Fuzhou, China), and visualized using a DAB kit (MXB Biotechnologies) according to the manufacturer’s instructions. The slides were counterstained with hematoxylin, dehydrated in a graded ethanol series, cleared in xylene, cover-slipped, and photographed under a light microscope (Axioscope 5; Carl Zeiss).

For immunofluorescent staining, the slides were washed three times with PBS to remove unbound primary antibodies and then incubated with goat anti-rabbit-Alexa Fluor™488 (A-11008; Thermo Fisher Scientific, Waltham, MA, USA) at 37 °C for 30 min, counterstained with 200 ng/mL 4′,6-diamidino-2-phenylindole (DAPI; Sigma-Aldrich) for 5 min to label nuclei, and imaged under a confocal microscope (LEICA DMi8; Leica Microsystems, Wetzlar, Germany).

### Terminal deoxynucleotidyl transferase dUTP nick end labeling (TUNEL) assay

To analyze apoptosis in mouse lung tissues, hydrated lung sections were evaluated using a TUNEL assay kit (Beyotime Biotechnology, Shanghai, China) per the manufacturer’s instructions and photographed under a light microscope (Axioscope 5; Carl Zeiss).

### DNA and siRNA transfection

The cells were transfected with plasmids and siRNAs using Lipofectamine 2000 (Thermo Fisher Scientific) according to the manufacturer’s instructions. The plasmids and siRNAs used are listed in Extended Data Tables 1 and 2.

### Lentiviral preparation and infection

To prepare lentiviruses, HEK293T cells were transiently transfected with pLVX-EF1alpha-2xStrep-IRES-Puro (control plasmid) or pLVX-EF1alpha-SARS-CoV-2-N-2xStrep-IRES-Puro (N protein-expressing plasmid), together with pMD2.G (Addgene, Cambridge, MA, USA) and psPAX2 (Addgene) using Lipofectamine 2000 according to the manufacturer’s recommendations. Seventy-two hours post-transfection, the supernatant was passed through a 0.45-µm syringe filter unit (EMD Millipore, Burlington, MA, USA). HBLV-EGFP-LC3-PURO lentivirus was purchased from Hanbio Biotechnology (Shanghai, China).

For infection, A549, BEAS-2B, and HEK293T cells at approximately 50% confluency were infected with lentivirus (multiplicity of infection [MOI] = 0.3) in the presence of 4 μg/mL polybrene (Sigma-Aldrich). Forty-eight hours post-infection, the cells were cultured in the presence of 2 μg/mL puromycin to remove uninfected cells. The cells infected with the control lentivirus were designated as A549-Control (A549-Ctrl), BEAS-2B-Control (BEAS-2B-Ctrl), and HEK293T-Control (HEK293T-Ctrl). The cells infected with the N protein-expressing lentivirus were designated as A549-N protein (A549-N), BEAS-2B-N protein (BEAS-2B-N), and HEK293T-N protein (HEK293T-N).

### Comet assay

Comet assay was performed as previously described^24^. Briefly, cells were washed, trypsinized, and suspended in 37 °C PBS at a concentration of 2 × 10^5^ cells/mL. A low-melting-point agarose (0.7% [m/v]; Solarbio, Beijing, China) suspension at 37 °C was then added to the cells at a 4:1 volume ratio and immediately transferred to a slide pre-coated with 0.6% (m/v) regular agarose (Solarbio). The slides were cover-slipped and incubated for 10 min at 4 °C. Subsequently, the coverslip was gently removed, and the slide was immersed in prechilled lysis solution (2.5 M NaCl, 100 mM ethylenediamine tetraacetic acid [EDTA], 10 mM Tris-HCl with 10% [v/v] DMSO, and 1% [v/v] Triton X-100 freshly added, pH 10.0) for 90 min at 4 °C. After lysis, the slides were washed with prechilled electrophoretic solution (1 mM EDTA, 300 mM NaOH, pH > 13.0), incubated in electrophoretic solution for 30 min at 4 °C, and subjected to electrophoresis at 25 V for 30 min at 4 °C in the dark. The slides were neutralized with 400 mM Tris (pH 7.5) for 30 min at 4 °C, dehydrated with 75% (v/v), 95% (v/v), and absolute ethanol for 5 min each at 25 °C, and stained with 0.1 μg/mL ethidium bromide (Sigma-Aldrich) for 30 min in the dark. Comet images were visualized using a fluorescence microscope (LEICA DMI3000 B; Leica Microsystems) and analyzed using the CASP software v1.2.3b2 (CaspLab, Wroclaw, Poland).

### Cytoplasmic/nuclear RNA extraction

Cytosolic and nuclear RNAs were extracted using a Cytoplasmic & Nuclear RNA Purification Kit (Norgen Biotek, Thorold, ON, USA) according to the manufacturer’s protocol.

### Luciferase reporter assay

To examine the effect of SARS-CoV-2 on RNA silencing, HEK293T-hACE2 cells were prepared by transfecting the pLVX-3×FLAG-hACE2 plasmid into HEK293T cells, seeded in a 48-well plate (2 × 10^4^ cells per well) 24 h before transfection, and co-transfected with 20 ng pRL-CMV (Promega, Madison, WI, USA), 100 ng pGL3-Control vector (Promega), and 100 ng pLKO.1-shNC-puro (shNC) or pLKO.1-shFluc-puro (shFluc). The cells were then infected with SARS-CoV-2 24 h post-transfection, and luciferase assays were performed 24 h post-infection.

To examine the effect of SARS-CoV-2 on RNA splicing, HEK293T-hACE2 cells were co-transfected in a 48-well plate with 20 ng pRL-CMV (Promega) and 100 ng CMV-LUC2CP/ARE (Negative-luc) or CMV-LUC2CP/intron/ARE (Intron-luc). The cells were then infected with SARS-CoV-2 24 h post-transfection, and luciferase assays were performed 24 h post-infection.

To assess the effect of SARS-CoV-2-encoding proteins on RNA silencing, HEK293T cells were co-transfected in a 48-well plate with 20 ng pRL-CMV, 100 ng pGL3-Control vector (Promega), and 100 ng shNC or shFluc, together with pLVX-EF1alpha-2xStrep-IRES-Puro (Ctrl), pLVX-EF1alpha-SARS-CoV-2-S-2xStrep-IRES-Puro (S), pLVX-EF1alpha-SARS-CoV-2-N-2xStrep-IRES-Puro (N), pLVX-EF1alpha-SARS-CoV-2-M-2xStrep-IRES-Puro (M), or pLVX-EF1alpha-SARS-CoV-2-E-2xStrep-IRES-Puro (E); luciferase assays were performed 48 h post-transfection.

To assess the effect of SARS-CoV-2-encoding proteins on RNA splicing, HEK293T cells were co-transfected in a 48-well plate with 20 ng pRL-CMV and 100 ng Negative-luc or Intron-luc, together with plasmids encoding S, N, M, or E; luciferase assays were performed 48 h post-transfection.

To assess the effect of N protein on RNA silencing, HEK293T-Ctrl and HEK293T-N cells were co-transfected in a 48-well plate with 20 ng pRL-CMV (Promega), 100 ng pGL3-Control vector (Promega), and 100 ng shNC or shFluc; luciferase assays were performed 48 h post-transfection.

To examine the effect of N protein on RNA splicing, HEK293T-Ctrl and HEK293T-N cells in a 48-well plate were co-transfected with 20 ng pRL-CMV (Promega) and 100 ng Negative-luc or Intron-luc; luciferase assays were performed 48 h post-transfection.

To examine whether SRSF3 and hnRNPA3 overexpression can rescue the inhibitory effect of N protein on RNA splicing, HEK293T-Ctrl or HEK293T-N cells in were co-transfected in a 48-well plate with 20 ng pRL-CMV (Promega) and 100 ng Negative-luc or Intron-luc, together with pCDH-CMV-MCS-EF1-Puro-SRSF3(h) (SRSF3) or pCDH-CMV-MCS-EF1-Puro-hnRNPA3(h) (hnRNPA3); luciferase assays were performed 48 h post-transfection.

To evaluate the effect of N protein on proteotoxic stress, A549-Ctrl or A549-N cells were co-transfected in a 48-well plate with 20 ng pRL-CMV (Promega) and 200 ng pCI-neo Fluc-EGFP or pCI-neo FlucDM-EGFP; luciferase assays were performed 48 h post-transfection.

Luciferase activity was measured using a dual-luciferase reporter assay system (Promega) according to the manufacturer’s protocol. pRL-CMV expressing *Renilla* luciferase under the *CMV* promoter was used as an external control to correct for differences in transfection and harvest efficiencies.

### Immunoblotting

Cells and tissues were lysed in radioimmunoprecipitation assay buffer supplemented with protease inhibitor cocktail tablets (Roche, Basel, Switzerland) and phosphatase inhibitor cocktail (APExBIO Technology LLC, Houston, TX, USA) at 4 °C; the protein concentration was measured using a Bicinchoninic Acid (BCA) Protein Assay Kit (Beyotime Biotechnology). Equal amounts of proteins were resolved by sodium dodecyl sulfate (SDS) polyacrylamide gel electrophoresis and transferred to a polyvinylidene fluoride membrane (Bio-Rad Laboratories, Hercules, CA, USA). The protein amount on the membrane was detected using Ponceau S Staining Solution (Thermo Fisher Scientific). Membranes were blocked with 5% bovine serum albumin in TBST (20 mM Tris-HCl [pH 7.5], 150 mM NaCl, and 0.1% Tween 20) for 1 h and subsequently probed with the indicated primary antibodies listed in Extended Data Table 3, followed by incubation with appropriate horseradish peroxidase-conjugated secondary antibodies (Extended Data Table 3). The blots were then treated with Clarity Western Enhanced Chemiluminescence Substrate (Bio-Rad Laboratories) and exposed to X-ray films (FujiFilm, Tokyo, Japan) in a dark room or visualized using the ChampChemi imaging system (Beijing Sage Creation Science, Beijing, China).

### Reverse transcription-quantitative polymerase chain reaction (RT-qPCR) analysis

Total RNA was prepared using TRIzol reagent (Life Technologies, Waltham, MA, USA) and then subjected to gel electrophoresis to assess RNA quality. The purified RNA was reverse-transcribed using the HiScript III RT SuperMix for qPCR (+gDNA wiper) kit (Vazyme Biotech Co., Ltd., Nanjing, China) for mRNAs and the miDETECT A Track miRNA qRT-PCR Starter Kit (RiboBio, Guangzhou, China) for miRNAs and pre-miRNAs. SYBR Green real-time PCR was performed using the ChamQ Universal SYBR qPCR Master Mix (Vazyme Biotech Co., Ltd.) and the ABI 7500 FAST sequence detection system (Life Technologies). The expression levels in all samples were normalized to those of *GAPDH* for mRNAs and *U6* for miRNAs and pre-miRNAs unless otherwise noted. Primer sequences are listed in Extended Data Table 4.

### Annexin V-fluoresceine isothiocyanate (FITC)/propidium iodide apoptosis assay

Cell apoptosis was detected via flow cytometry using an FITC Annexin V Apoptosis Detection Kit (BD Biosciences, San Jose, CA, USA), according to the manufacturer’s instructions.

### Immunoprecipitation (IP) assay

The cells were lysed with IP binding buffer (20 mM Tris [pH 7.5], 150 mM NaCl, and 1% [v/v] Triton X-100) supplemented with protease inhibitor cocktail tablets (Roche) at 4 °C for 30 min with continuous rotation, followed by centrifugation at 13,000 × *g* for 10 min. Equal amounts of the lysate were incubated overnight at 4 °C with anti-nucleocapsid protein antibody (40588-RC02; Sino Biological, Beijing, China), anti-p62 (SQSTM1) antibody (18420-1-AP; Proteintech), or normal rabbit immunoglobulin G (IgG; A7058; Beyotime Biotechnology) and precipitated with Protein A/G magnetic beads (Selleck Chemicals, Houston, TX, USA). The beads were washed five times with IP washing buffer (50 mM Tris [pH 7.5], 150 mM NaCl, 1% [v/v] Triton X-100), supplemented with protease inhibitor cocktail tablets (Roche), and eluted with SDS loading buffer at 100 °C for 10 min. The precipitated immune complexes were subjected to immunoblotting analysis to detect the interacting proteins.

To determine whether inhibiting the RNA-binding activity of N protein disrupted the interaction between N protein and Dicer, XPO5, SRSF3, or hnRNPA3, cells were treated with PJ34 (50 μM) for 2 h before being lysed.

To determine whether RNA is involved in the interaction between N protein and Dicer, XPO5, SRSF3, or hnRNPA3, the cellular extracts were treated with RNase A (1 mg/mL; TakaRa, Dalian, China), RNase T1 (20 U/mL; Thermo Fisher Scientific), RNase V1 (20 U/mL; Life Technologies), and RNase I (20 U/mL; Life Technologies) for 15 min at 37 °C before IP.

### RNA-immunoprecipitation (RIP) assay

The cells were cross-linked with 1% (v/v) formaldehyde. Crosslinking was stopped by adding 200 mM glycine, and the cells were lysed in 0.5 mL of ice-cold RIP lysis buffer (50 mM Tris-HCl [pH 8.0], 150 mM KCl, 5 mM EDTA, 0.1% [m/v] SDS, 1% [v/v] Triton X-100, 0.5% [m/v] sodium deoxycholate, and 0.5 mM dithiothreitol [DTT]) supplemented with 500 U/mL RNase inhibitor (Vazyme Biotech Co., Ltd) and protease inhibitor cocktail tablets (Roche) and sonicated at 4 °C for 4 min (30 s on and 30 s off) on a QSonica Q800R3 sonicator (QSonica, Newton, CT, USA) under 30% power. The cell lysate was centrifuged at 12,000 × *g* and 4 °C for 10 min. The supernatants were mixed with equal volumes of RIP binding buffer (25 mM Tris-HCl [pH 7.5], 150 mM KCl, 5 mM EDTA, 0.5% [v/v] NP-40, and 0.5 mM DTT) supplemented with 500 U/mL RNase inhibitor (Vazyme Biotech Co., Ltd) and protease inhibitor cocktail tablets (Roche) and incubated with an anti-nucleocapsid protein antibody (Sino Biological) or normal rabbit IgG control (Beyotime Biotechnology) at 4 °C for 6 h. Subsequently, they were incubated with Protein A/G magnetic beads at 4 °C for 4 h. The beads were then washed five times using RIP washing buffer (50 mM Tris-HCl [pH 7.5], 150 mM KCl, 5 mM EDTA, 0.5% [v/v] NP-40, and 0.5 mM DTT) supplemented with 500 U/mL RNase inhibitor (Vazyme Biotech Co., Ltd.) and protease inhibitor cocktail tablets (Roche); RNA was extracted using the TRIzol reagent and quantified via RT-qPCR. Relative enrichment was calculated as the ratio between specific antibodies and the normal IgG control, defined as 1.

### Protein synthesis measurement

Nascent polypeptides were labeled by treating cells with 10 μg/mL puromycin (MedChemExpress) for 1 h, followed by immunoblotting with an anti-puromycin antibody (MABE343; Sigma-Aldrich, 1:5000).

### Soluble and insoluble protein isolation

Ultracentrifugation was used to isolate soluble and insoluble proteins, as previously described^61^. Briefly, the cells were lysed with Triton X-100 buffer (1% [v/v] Triton X-100, 150 mM NaCl, 50 mM Tris-HCl [pH 8.0], and 1 mM EDTA) supplemented with protease inhibitor cocktail (Roche) and sonicated at 4 °C for 1 min (10 s on and 10 s off) on a QSonica Q800R3 sonicator (QSonica). The protein concentration was quantified using a BCA Protein Assay Kit (Beyotime Biotechnology) to ensure that the starting amounts of the protein samples were equal. The cell lysate was centrifuged at 100,000 × *g* for 30 min using a Beckman TL-100 Ultracentrifuge (Beckman, Palo Alto, CA, USA) at 4 °C. The soluble proteins in the supernatant were collected for immunoblotting. The insoluble pellet was re-suspended in SDS lysis buffer (Triton X-100 buffer, 2% [m/v] SDS), supplemented with protease inhibitor cocktail tablets (Roche), and sonicated on a QSonica Q800R3 sonicator (QSonica) until the pellet became invisible. The BCA assay was used to quantify the protein concentration in the pellet fraction. Soluble and insoluble protein samples were analyzed using immunoblotting.

### Confocal microscopy

#### Immunofluorescence

Immunofluorescence staining was used to detect cytosolic dsDNA, γ-H2AX foci, the R-loop, and co-localization of N protein with p62. The cells were fixed with 4% (m/v) paraformaldehyde diluted in PBS and incubated with anti-dsDNA (Sigma-Aldrich, 1:500), anti-γ-H2AX (2577; Cell Signaling Technology, Danvers, MA, USA, 1:500), anti-DNA-RNA hybrid antibody, clone S9.6 (MABE1095; Sigma-Aldrich, 1:500), anti-nucleocapsid protein (Sino Biological, 1:100), or anti-p62 (88588; Cell Signaling Technology, 1:200) antibody at 4 °C overnight. The cells were then incubated with goat anti-mouse-Alexa Fluor™ 488 (A-11001; Thermo Fisher Scientific, 1:1000) for dsDNA, R-loop, and p62; with goat anti-rabbit-Alexa Fluor™ 488 (Thermo Fisher Scientific, 1:500) for γ-H2AX; or with goat anti-rabbit-Alexa Fluor™ 555 (A-21428; Thermo Fisher Scientific, 1:500) for N protein at 37 °C for 30 min. Subsequently, the nuclei were counterstained with 200 ng/mL DAPI (Sigma-Aldrich) for 5 min to label nuclei, followed by imaging using a confocal microscope (LEICA DMi8; Leica Microsystems). To calculate the average number of γ-H2AX foci or R-loop foci per nucleus, a minimum of 200 cells were counted in at least three independent biological replicate experiments.

#### LC3 puncta imaging

A549 cells were infected with HBLV-EGFP-LC3-PURO lentivirus (Hanbio Biotechnology, Shanghai, China); transduced cells were selected using 2 µg/mL puromycin (MedChemExpress). EGFP-LC3-expressing cells were transfected with an N protein-expressing plasmid or a control plasmid. The cells were fixed with 4% (m/v) paraformaldehyde 48 h post-transfection, followed by counterstaining with 200 ng/mL DAPI (Sigma-Aldrich) for 5 min, and imaged using laser scanning confocal microscopy. To calculate the average number of EGFP-LC3-positive puncta per cell, a minimum of 200 cells were counted in at least three independent experiments.

#### Proteotoxic stress analysis

A549-Ctrl and A549-N cells were transfected with pCI-neo Fluc-EGFP or pCI-neo FlucDM-EGFP and fixed with 4% (m/v) paraformaldehyde 48 h post-transfection, followed by counterstaining with 200 ng/mL DAPI (Sigma-Aldrich) for 5 min, and imaged using laser scanning confocal microscopy. To calculate the average number of Fluc-EGFP aggregates per cell, a minimum of 200 cells were counted in at least three independent experiments.

### Gene Expression Omnibus (GEO) dataset analysis

Transcriptome data were downloaded from the GEO database (https://www.ncbi.nlm.nih.gov/gds/). The GSE178824 dataset comprised RNA sequencing data of monocytic myeloid-derived suppressor cells from patients with severe or asymptomatic COVID-19; GSE159585 dataset comprised RNA sequencing data for lung tissues from deceased patients with COVID-19 and individuals without COVID-19; and GSE209891 dataset comprised RNA sequencing data of lung tissues from mice of different ages. Differentially expressed genes were identified using the limma (version 3.54.2) R package^62^. The genes of the miRNA processing and RNA splicing pathways were retrieved from the biological process categories in the Molecular Signatures Database (https://www.gsea-msigdb.org/gsea/msigdb). The Z-score of each gene was calculated for each biological replicate and plotted and clustered using the heatmap function in R v 4.1.0 (R Foundation for Statistical Computing, Vienna, Austria) with hierarchical clustering based on the Euclidean distance.

### In silico protein–protein interaction analysis

Putative protein interactors of N protein were retrieved from the Biological General Repository for Interaction Datasets (https://thebiogrid.org/)^38^. Pathway enrichment of putative N protein interactors was performed using the Metascape webserver (http://metascape.org)^63^.

### Small RNA-seq analysis

Small RNA sequencing was performed by Biomarker Technologies Corporation (Beijing, China). The clean sequencing reads were aligned to the Silva database, GtRNAdb database, Rfam database, and Repbase database using Bowtie 1.0.0 (https://bowtie-bio.sourceforge.net/index.shtml) to filter small RNAs derived from ribosomal RNA, transfer RNA, small nuclear RNA, small nucleolar RNA, other noncoding RNAs, and repeat sequences. The remaining reads were aligned to miRbase to identify known miRNAs. Differentially expressed miRNAs (∣fold change∣ > 1.5, and p < 0.05) were identified using the DESeq (version 1.10.1) R package^64^. To generate a heatmap of the top 30 differentially expressed miRNAs with the highest expression, the Z-score of each miRNA was calculated for each biological replicate and then plotted and clustered using the heatmap function in R v 4.1.0 (R Foundation for Statistical Computing), with hierarchical clustering based on the Euclidean distance.

### Alternative splicing analysis

Full-length transcriptome sequencing was performed by Biomarker Technologies Corporation using Oxford Nanopore Technology^65^. Alternative splicing events were analyzed using the PSI-Sigma method, as previously described^66^. ΔPSI was calculated as follows: ΔPSI = (PSI value of A549-N cells) – (PSI value of A549-Ctrl cells). An absolute value of ΔPSI ≥ 10% was set as the cutoff for alternative splicing events with significant changes^67^.

To analyze the overall intron retention (IR) levels in A549-Ctrl and A549-N cells, the full-length transcriptome sequencing data were mapped to the human genome (hg19; known reference genes from the University of California, Santa Cruz [UCSC]) using Minimap2^68^. Exons were annotated using RefSeq, and introns were defined as the genomic regions not covered by any exons annotated using RefSeq^69^. The fraction of transcripts with IR over total sequencing reads was calculated as follows: Fraction of transcripts with IR = isoform reads containing introns ÷ total sequencing reads. The relative fraction of transcripts with IR in A549-Ctrl cells was defined as 1.

The intronic and exonic RNA levels of *GPX1*, *TERT*, and *CCN1* in A549-Ctrl or A549-N cells and BEAS-2B-Ctrl or BEAS-2B-N cells were determined via RT-qPCR. The relative ratio of intronic and exonic RNA levels was calculated and defined as 1 in Ctrl cells.

### Quantification and statistical analysis

The microscopy image quantification was performed using Fiji 2.3.0^70^. Data were presented as mean ± standard deviation of at least three independent experiments. Normal distribution was confirmed using the Shapiro–Wilk normality test, and homogeneity of variance was tested using Levene’s test. Student’s *t*-test was used to compare the differences between two groups. In all experiments, the significance level was set as α = 0.05, and p < 0.05 indicated significant intergroup differences. Statistical analyses were performed using SPSS Statistics 23.0 (IBM SPSS Statistics, IBM Corporation, Armonk, NY, USA). Additional information regarding statistical tests, p values, and sample size are described in the figure legends.

### Reporting summary

Further information on research design is available in the Nature Portfolio Reporting Summary linked to this article.

### Data availability

All data needed to evaluate the conclusions are included in the paper and/or the Supplemental Information. The small RNA and full-length transcriptome sequencing data were deposited in the NCBI Sequence Read Archive under BioSample accession numbers PRJNA1000280 and PRJNA1000513, respectively.

## Notes

### Competing Interest Statement

The authors have declared no competing interest.

