## Supplemental information for "SARS-CoV-2 N protein-induced Dicer, XPO5, SRSF3, and hnRNPA3 downregulation causes pneumonia"

### Inventory of Supporting Information

**This PDF file includes:**

Extended Data Figures 1 to 7

Extended Data Tables 1 to 3

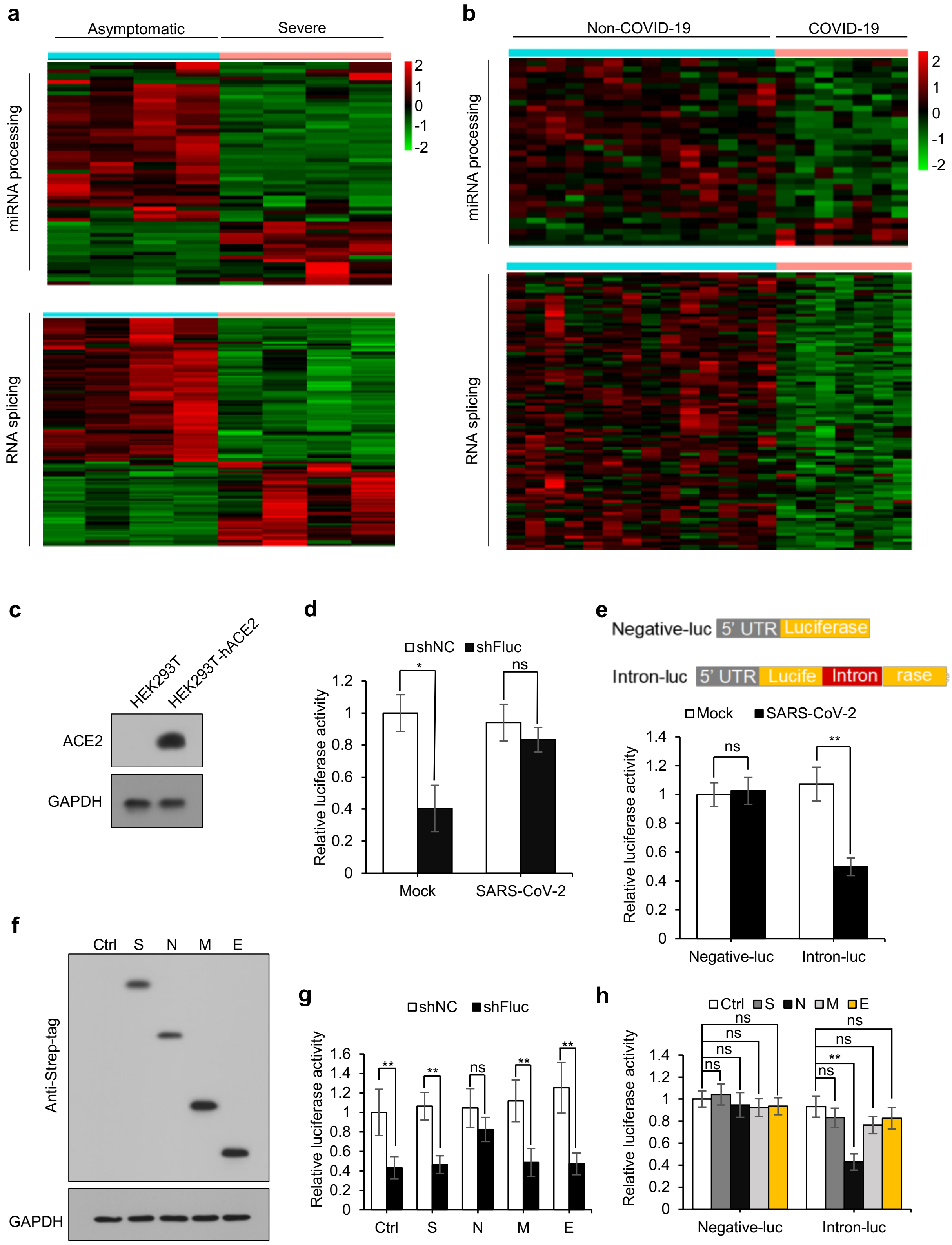
a
Asymptomatic
Severe
miRNA processing
RNA splicing
2
1
0
-1
-2
b
Non-COVID-19
COVID-19
miRNA processing
RNA splicing
c
HEK293T
HEK293T-hACE2
ACE2
GAPDH
d
shNC
shFluc
Relative luciferase activity
Mock
SARS-CoV-2
e
Negative-luc
5' UTR
Luciferase
Intron-luc
Lucife
Intron
rase
Mock
SARS-CoV-2
Negative-luc
Intron-luc
f
Ctrl
S
N
M
E
Anti-Strep-tag
GAPDH
g
shNC
shFluc
Relative luciferase activity
Ctrl
S
N
M
E
h
Ctrl
S
N
M
E
Relative luciferase activity
Negative-luc
Intron-luc

**Extended Data Figure 1. SARS-CoV-2 N protein suppresses RNAi and RNA splicing;** related to Fig. 1. **(a)** Heatmap depicting the mRNA levels of RNAi components and RNA splicing factors in monocytic-myeloid-derived suppressor cells (M-MDSCs) from patients with severe or asymptomatic coronavirus disease 2019 (COVID-19). The RNA sequencing data were downloaded from the Gene Expression Omnibus (GEO) database (GSE178824). **(b)** Heatmap depicting the mRNA levels of RNAi components and RNA splicing factors in lung tissues from deceased patients with COVID-19 and individuals without COVID-19. The RNA sequencing data were downloaded from the GEO database (GSE159585). **(c)** Ectopic expression of hACE2 in HEK293T cells was confirmed via immunoblotting analysis. **(d)** Effect of SARS-CoV-2 on RNAi. HEK293T-hACE2 cells were co-transfected with plasmids encoding firefly luciferase (pGL3-Control) and firefly luciferase-specific shRNA (shFluc) or control shRNA (shNC) and then infected with SARS-CoV-2 or mock-infected 24 h post-transfection. Luciferase assay was performed 24 h post-infection. **(e)** Effect of SARS-CoV-2 on RNA splicing; upper: schematic of RNA splicing reporters; lower: HEK293T-hACE2 cells were transfected with the splicing reporter and infected with SARS-CoV-2 or mock-infected 24 h post-transfection. Luciferase assays were performed 24 h post-infection. **(f, g)** HEK293T cells co-transfected with pGL3-Control, shFluc, or shNC, and a panel of plasmids expressing different strep-tagged viral proteins. Immunoblotting with the anti-strep-tag antibody (f) and luciferase assay (g) were performed 48 h post-transfection. **(h)** Splicing reporter plasmids were co-transfected with plasmids encoding different viral proteins into HEK293T cells; luciferase assay was performed 48 h post-transfection. Data in d, e, g, and h are expressed as mean ± standard deviation (SD) of three biological replicates. $^{**}p < 0.01$; $^{*}p < 0.05$; ns, not significant ($p > 0.05$; two-sided Student's *t*-test). Ctrl: control plasmid; N: N protein; S: spike protein; M: membrane protein; E: envelope protein.

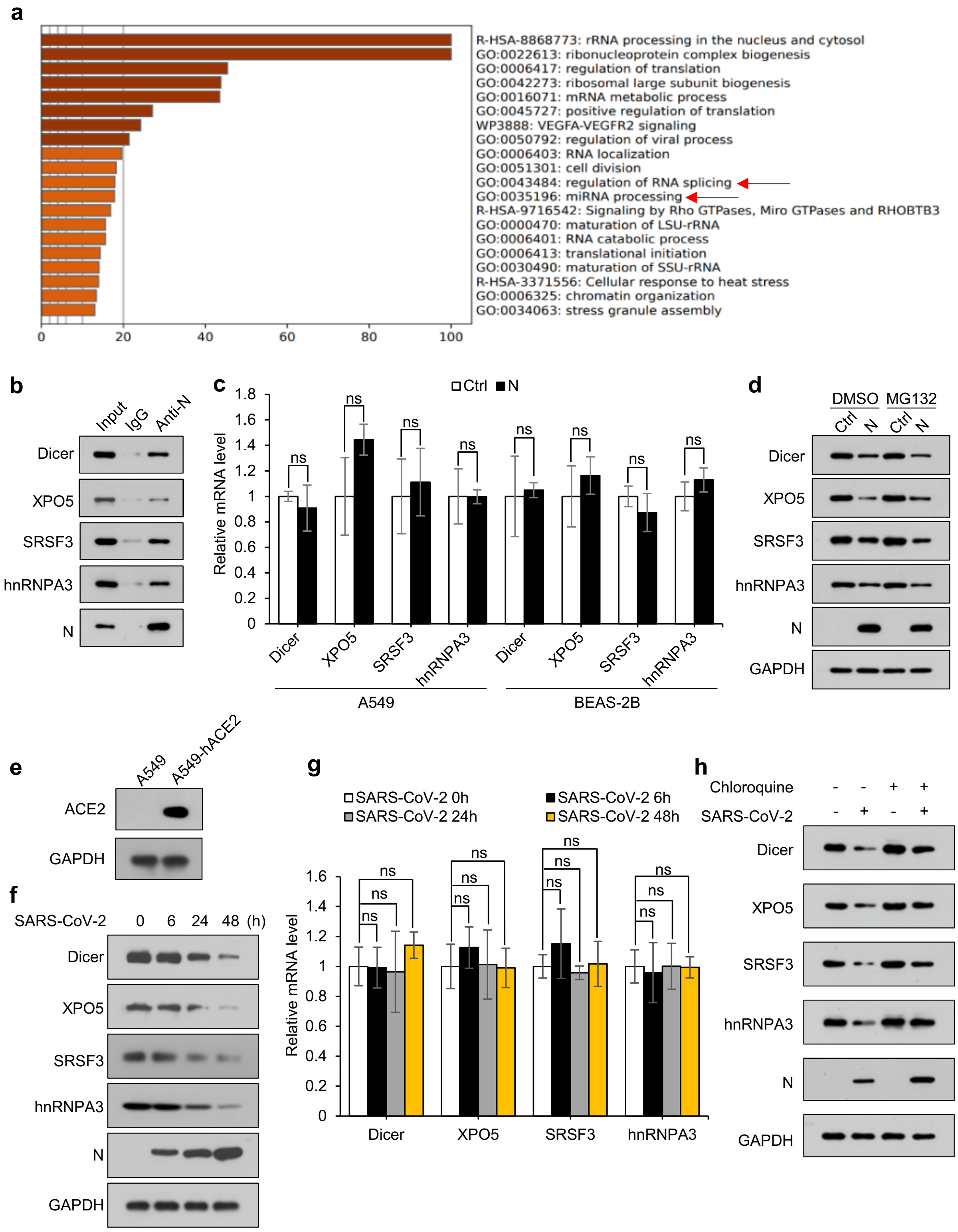
a
R-HSA-8868773: rRNA processing in the nucleus and cytosol
GO:0022613: ribonucleoprotein complex biogenesis
GO:0006417: regulation of translation
GO:0042273: ribosomal large subunit biogenesis
GO:0016071: mRNA metabolic process
GO:0045727: positive regulation of translation
WP3888: VEGFA-VEGFR2 signaling
GO:0050792: regulation of viral process
GO:0006403: RNA localization
GO:0051301: cell division
GO:0043484: regulation of RNA splicing
GO:0035196: miRNA processing
R-HSA-9716542: Signaling by Rho GTPases, Miro GTPases and RHOBTB3
GO:0000470: maturation of LSU-rRNA
GO:0006401: RNA catabolic process
GO:0006413: translational initiation
GO:0030490: maturation of SSU-rRNA
R-HSA-3371556: Cellular response to heat stress
GO:0006325: chromatin organization
GO:0034063: stress granule assembly
0 20 40 60 80 100
b
Input IgG Anti-N
Dicer
XPO5
SRSF3
hnRNPA3
N
c
Ctrl N
Relative mRNA level
1.8 1.6 1.4 1.2 1 0.8 0.6 0.4 0.2 0
ns
Dicer XPO5 SRSF3 hnRNPA3 Dicer XPO5 SRSF3 hnRNPA3
A549
BEAS-2B
d
DMSO MG132
Ctrl N Ctrl N
Dicer
XPO5
SRSF3
hnRNPA3
N
GAPDH
e
A549 A549-hACE2
ACE2
GAPDH
f
SARS-CoV-2 0 6 24 48 (h)
Dicer
XPO5
SRSF3
hnRNPA3
N
GAPDH
g
SARS-CoV-2 0h
SARS-CoV-2 6h
SARS-CoV-2 24h
SARS-CoV-2 48h
Relative mRNA level
1.6 1.4 1.2 1 0.8 0.6 0.4 0.2 0
ns
Dicer XPO5 SRSF3 hnRNPA3
h
Chloroquine - - + +
SARS-CoV-2 - + - +
Dicer
XPO5
SRSF3
hnRNPA3
N
GAPDH

**Extended Data Figure 2. SARS-CoV-2 N protein interacts with and induces the autophagic degradation of Dicer, XPO5, SRSF3, and hnRNPA3;** related to Fig. 1. **(a)** Metascape pathway enrichment of putative N protein interactors. **(b)** Lysates from BEAS-2B cells stably expressing N protein (BEAS-2B-N) were immunoprecipitated using anti-N protein antibody and immunoblotted with the indicated antibodies. **(c)** mRNA levels of *Dicer, XPO5, SRSF3*, and *hnRNPA3* in A549 cells stably transfected with control plasmid (A549-Ctrl), A549 cells stably expressing N protein (A549-N), and BEAS-2B cells stably transfected with control plasmid (BEAS-2B-Ctrl) and BEAS-2B-N cells. **(d)** Immunoblotting of the indicated proteins in A549-Ctrl or A549-N cells treated with or without a proteasome inhibitor MG132. **(e)** Ectopic expression of hACE2 in A549 cells confirmed via immunoblotting analysis. **(f, g)** A549-hACE2 cells were infected with SARS-CoV-2 or mock-infected; the protein (f) and mRNA (g) levels of Dicer, XPO5, SRSF3, and hnRNPA3 were determined at the three indicated time points after infection (g). **(h)** Immunoblotting of the indicated proteins in A549-hACE2 cells infected with SARS-CoV-2 or mock-infected and treated with or without chloroquine. Data in c and g are expressed as mean ± SD of three biological replicates. ns, not significant ($p > 0.05$) (two-sided Student's *t*-test). Ctrl: control plasmid; N: N protein; anti-N: anti-N protein antibody.

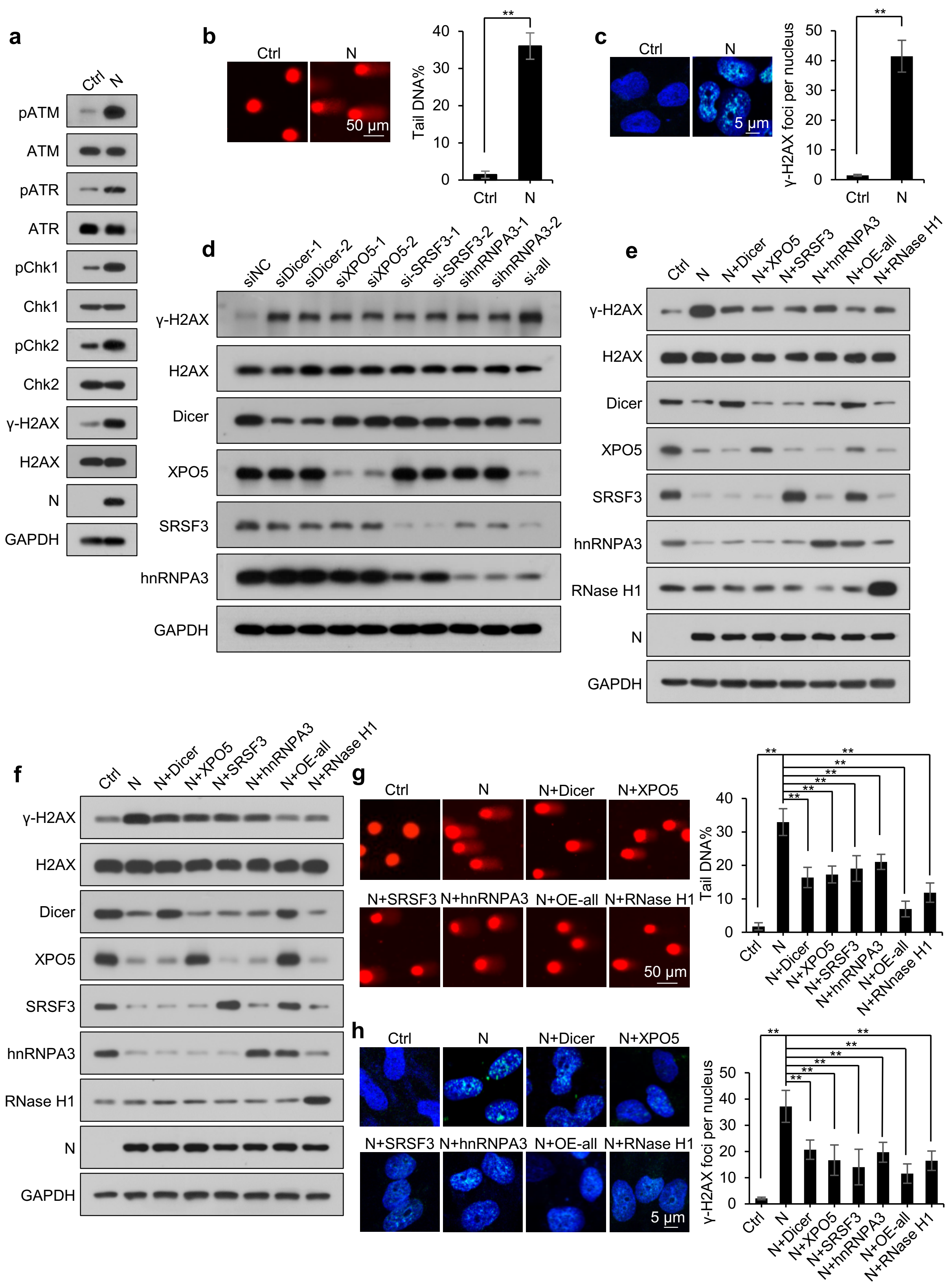

a
Ctrl
N
pATM
ATM
pATR
ATR
pChk1
Chk1
pChk2
Chk2
γ-H2AX
H2AX
N
GAPDH
b
Ctrl
N
50 μm
Tail DNA%
**
40
30
20
10
0
Ctrl
N
c
Ctrl
N
5 μm
γ-H2AX foci per nucleus
**
50
40
30
20
10
0
Ctrl
N
d
siNC
siDicer-1
siDicer-2
siXPO5-1
siXPO5-2
si-SRSF3-1
si-SRSF3-2
sihnRNPA3-1
sihnRNPA3-2
si-all
γ-H2AX
H2AX
Dicer
XPO5
SRSF3
hnRNPA3
GAPDH
e
Ctrl
N
N+Dicer
N+XPO5
N+SRSF3
N+hnRNPA3
N+OE-all
N+RNase H1
γ-H2AX
H2AX
Dicer
XPO5
SRSF3
hnRNPA3
RNase H1
N
GAPDH
f
Ctrl
N
N+Dicer
N+XPO5
N+SRSF3
N+hnRNPA3
N+OE-all
N+RNase H1
γ-H2AX
H2AX
Dicer
XPO5
SRSF3
hnRNPA3
RNase H1
N
GAPDH
g
Ctrl
N
N+Dicer
N+XPO5
N+SRSF3
N+hnRNPA3
N+OE-all
N+RNase H1
50 μm
Tail DNA%
**
40
30
20
10
0
Ctrl
N
N+Dicer
N+XPO5
N+SRSF3
N+hnRNPA3
N+OE-all
N+RNase H1
h
Ctrl
N
N+Dicer
N+XPO5
N+SRSF3
N+hnRNPA3
N+OE-all
N+RNase H1
5 μm
γ-H2AX foci per nucleus
**
50
40
30
20
10
0
Ctrl
N
N+Dicer
N+XPO5
N+SRSF3
N+hnRNPA3
N+OE-all
N+RNase H1

**Extended Data Figure 3. SARS-CoV-2 N protein induces DNA damage by downregulating Dicer, XPO5, SRSF3, and hnRNPA3 expression;** related to Fig. 2. **(a–c)** DNA damage in BEAS-2B-Ctrl or BEAS-2B-N cells was determined via immunoblotting analysis of the phosphorylation levels of ATM, ATR, Chk1, Chk2, and H2AX (a), comet assay (b), and immunofluorescence staining with anti-γ-H2AX antibody (c). **(d)** A549 cells were transfected with the indicated siRNAs and subjected to immunoblotting. **(e)** A549-Ctrl and A549-N cells transfected with a control plasmid or plasmids overexpressing Dicer, XPO5, SRSF3, hnRNPA3, or RNase H1 were subjected to immunoblotting. **(f–h)** BEAS-2B-Ctrl and BEAS-2B-N cells transfected with a control plasmid or plasmids expressing Dicer, XPO5, SRSF3, hnRNPA3, or RNase H1 were subjected to immunoblotting (f), comet assay (g), and immunofluorescence staining with anti-γ-H2AX antibody (h). Data in b, c, g, and h are expressed as mean ± SD of three biological replicates ($n \geq 200$ cells). $^{**}p < 0.01$ (two-sided Student's *t*-test). Ctrl: control plasmid; N: N protein; siNC: negative control siRNA; si-all: cells transfected with siDicer, siXPO5, siSRSF3, and sihnRNPA3 together; OE-all: cells transfected with Dicer-, XPO5-, SRSF3-, and hnRNPA3-expressing plasmids together.

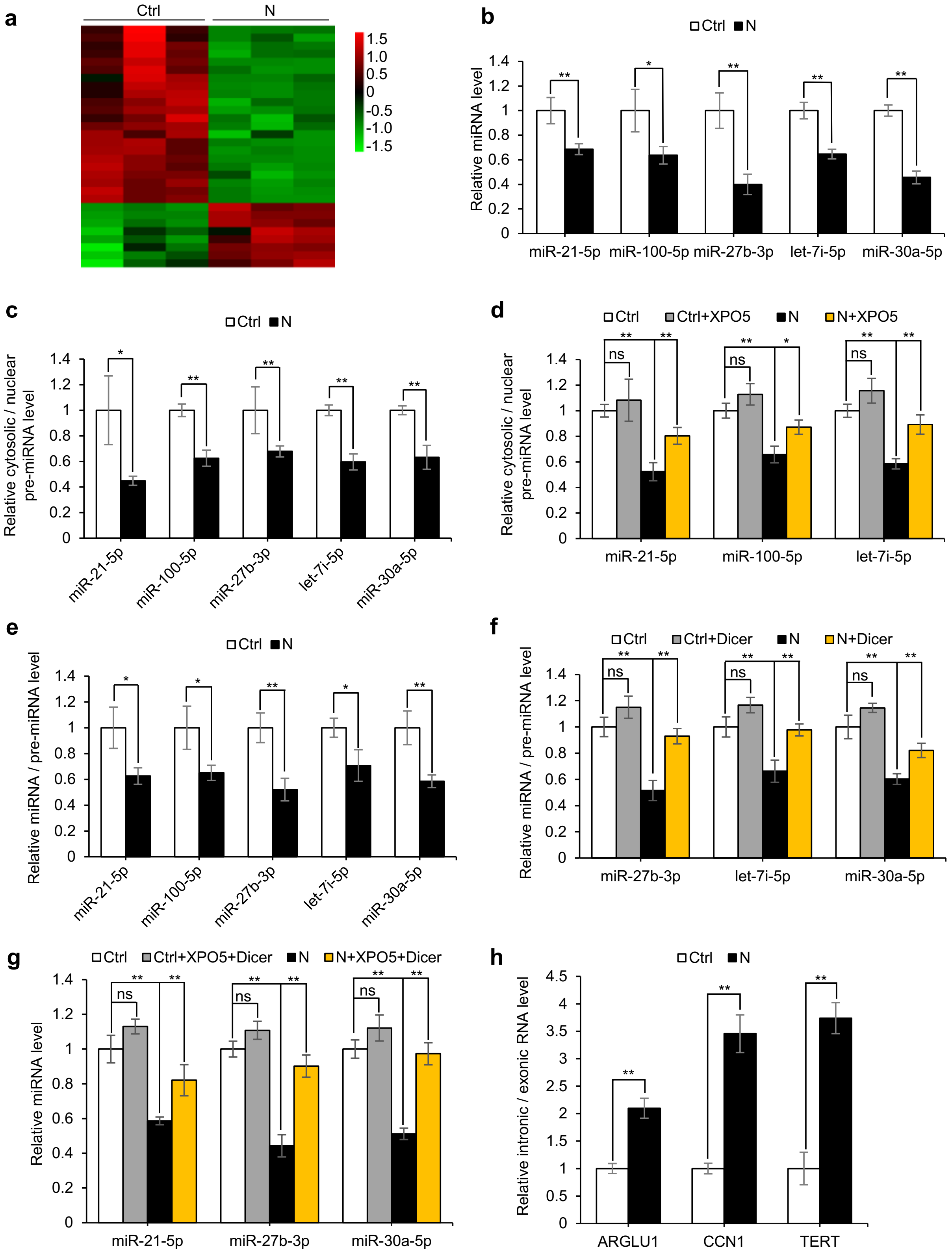

a
Ctrl
N
1.5
1.0
0.5
0
-0.5
-1.0
-1.5
b
Ctrl N
Relative miRNA level
miR-21-5p miR-100-5p miR-27b-3p let-7i-5p miR-30a-5p
c
Ctrl N
Relative cytosolic / nuclear pre-miRNA level
miR-21-5p miR-100-5p miR-27b-3p let-7i-5p miR-30a-5p
d
Ctrl Ctrl+XPO5 N N+XPO5
Relative cytosolic / nuclear pre-miRNA level
miR-21-5p miR-100-5p let-7i-5p
e
Ctrl N
Relative miRNA / pre-miRNA level
miR-21-5p miR-100-5p miR-27b-3p let-7i-5p miR-30a-5p
f
Ctrl Ctrl+Dicer N N+Dicer
Relative miRNA / pre-miRNA level
miR-27b-3p let-7i-5p miR-30a-5p
g
Ctrl Ctrl+XPO5+Dicer N N+XPO5+Dicer
Relative miRNA level
miR-21-5p miR-27b-3p miR-30a-5p
h
Ctrl N
Relative intronic / exonic RNA level
ARGLU1 CCN1 TERT

**Extended Data Figure 4. SARS-CoV-2 N protein represses miRNA biogenesis and splicing;** related to Fig. 3. **(a)** Heatmap of miRNA expression in BEAS-2B-Ctrl and BEAS-2B-N cells based on small RNA sequencing. **(b)** Quantification of five miRNAs in BEAS-2B-Ctrl and BEAS-2B-N cells. **(c)** Ratio of cytosolic pre-miRNA levels to nuclear pre-miRNA levels in BEAS-2B-Ctrl and BEAS-2B-N cells. **(d)** Ratio of cytosolic pre-miRNA levels to nuclear pre-miRNA levels in BEAS-2B-Ctrl and BEAS-2B-N cells transfected with a plasmid expressing XPO5 or control plasmid. **(e)** Ratio of mature miRNA levels to pre-miRNA levels in BEAS-2B-Ctrl and BEAS-2B-N cells. **(f)** Ratio of mature miRNA levels to pre-miRNA levels in BEAS-2B-Ctrl and BEAS-2B-N cells transfected with a Dicer-expressing or control plasmid. **(g)** miRNA levels in BEAS-2B-Ctrl and BEAS-2B-N cells transfected with XPO5-expressing and Dicer-expressing plasmids or a control plasmid. **(h)** Intronic and exonic RNA levels of the indicated genes in BEAS-2B-Ctrl and BEAS-2B-N cells. Data in b–h are expressed as mean ± SD of three biological replicates. **p < 0.01; *p < 0.05; ns, not significant (p > 0.05; two-sided Student's *t*-test). Ctrl: control plasmid; N: N protein.

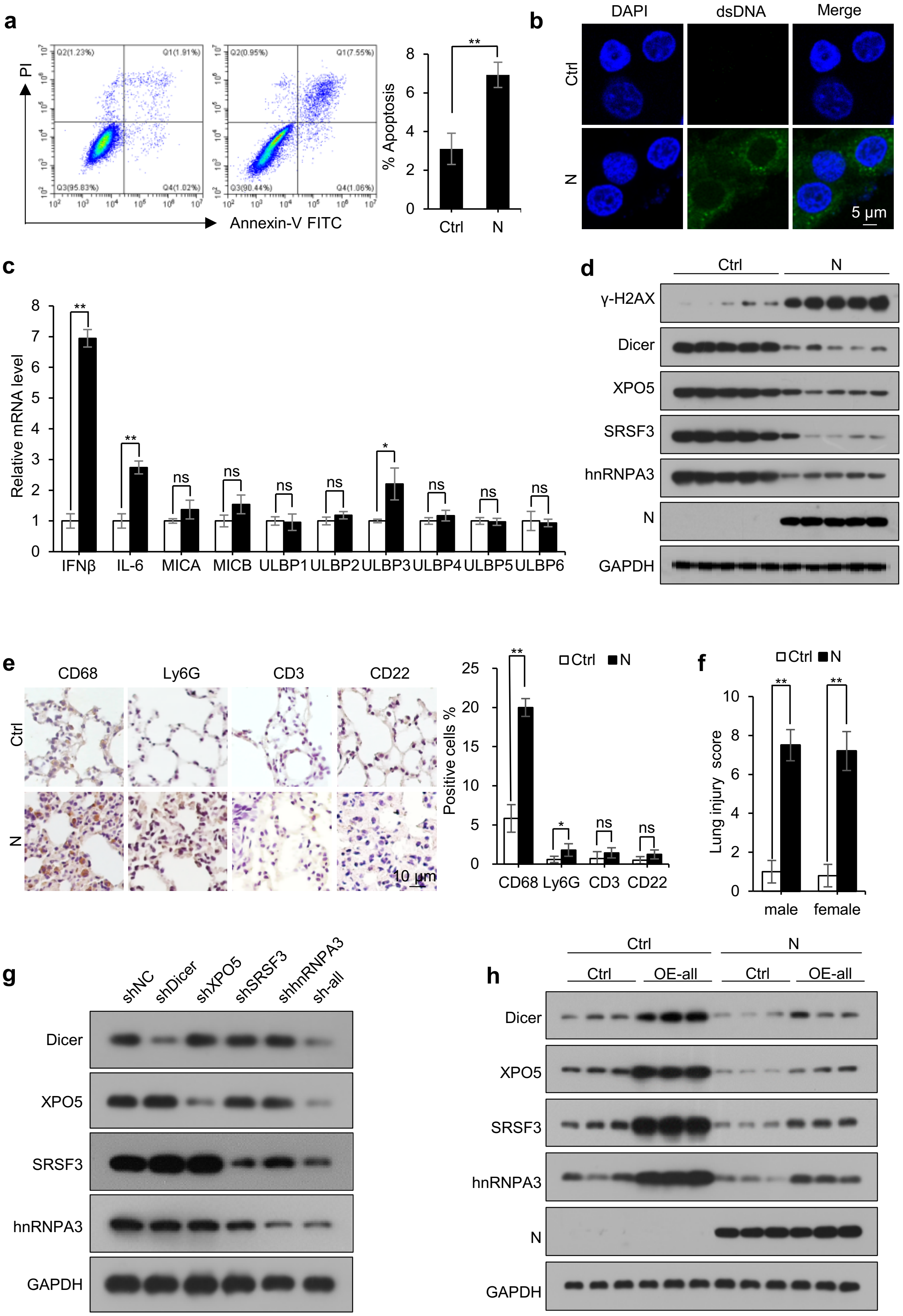
a
PI
Q2(1.23%)
Q1(1.91%)
Q3(95.83%)
Q4(1.02%)
Q2(0.95%)
Q1(7.55%)
Q3(90.44%)
Q4(1.06%)
Annexin-V FITC
% Apoptosis
**
Ctrl
N
b
DAPI
dsDNA
Merge
Ctrl
N
5 μm
c
Relative mRNA level
**
**
ns
ns
ns
ns
*
ns
ns
ns
IFNβ
IL-6
MICA
MICB
ULBP1
ULBP2
ULBP3
ULBP4
ULBP5
ULBP6
d
Ctrl
N
γ-H2AX
Dicer
XPO5
SRSF3
hnRNPA3
N
GAPDH
e
CD68
Ly6G
CD3
CD22
Ctrl
N
10 μm
Positive cells %
□Ctrl ■N
**
*
ns
ns
CD68
Ly6G
CD3
CD22
f
□Ctrl ■N
Lung injury score
**
**
male
female
g
shNC
shDicer
shXPO5
shSRSF3
shhnRNPA3
sh-all
Dicer
XPO5
SRSF3
hnRNPA3
GAPDH
h
Ctrl
N
Ctrl
OE-all
Ctrl
OE-all
Dicer
XPO5
SRSF3
hnRNPA3
N
GAPDH

**Extended Data Figure 5. SARS-CoV-2 N protein induces pneumonia by downregulating Dicer, XPO5, SRSF3, and hnRNPA3 expression;** related to Fig. 5. **(a)** Apoptosis in A549-Ctrl and A549-N cells measured using Annexin V-FITC and propidium iodide (PI) staining. **(b)** Representative confocal microscopy image of immunofluorescence detection of cytosolic DNA in A549-Ctrl or A549-N cells. **(c)** mRNA levels of *IFNβ, IL-6,* and *NKG2D* ligands in A549-Ctrl or A549-N cells. **(d, e)** Eight-week-old male mice were intranasally instilled with control- or N protein-expressing plasmid, and lung tissues were subjected to immunoblotting with the indicated antibodies (d), and immunohistochemical staining with antibodies against markers of different immune cells, including CD68 (macrophages), Ly6G (neutrophils), CD3 (T cells), and CD22 (B cells) (e). **(f)** Eight-week-old male or female mice were intranasally instilled with control- or N protein-expressing plasmid; lung tissues were subjected to HE staining to assess lung injury. **(g)** Immunoblotting of the indicated proteins in the lung tissues of mice intranasally instilled with plasmids expressing shNC or shDicer, shXPO5, shSRSF3, and shhnRNPA3 alone or in combination. **(h)** Immunoblotting of the indicated proteins in the lung tissues of mice intranasally instilled with N protein-expressing plasmid or co-instilled with N protein-expressing plasmid and plasmids expressing Dicer, XPO5, SRSF3, and hnRNPA3. Data in a and c are expressed as mean ± SD of three biological replicates. $^{**}p < 0.01$; $^{*}p < 0.05$; ns, not significant ($p > 0.05$; two-sided Student's *t*-test). Ctrl: control plasmid; N: N protein; shNC: negative control shRNA; sh-all: mice instilled with shDicer, shXPO5, shSRSF3, and shhnRNPA3 together; OE-all: mice instilled with Dicer-, XPO5-, SRSF3-, and hnRNPA3-expressing plasmids together.

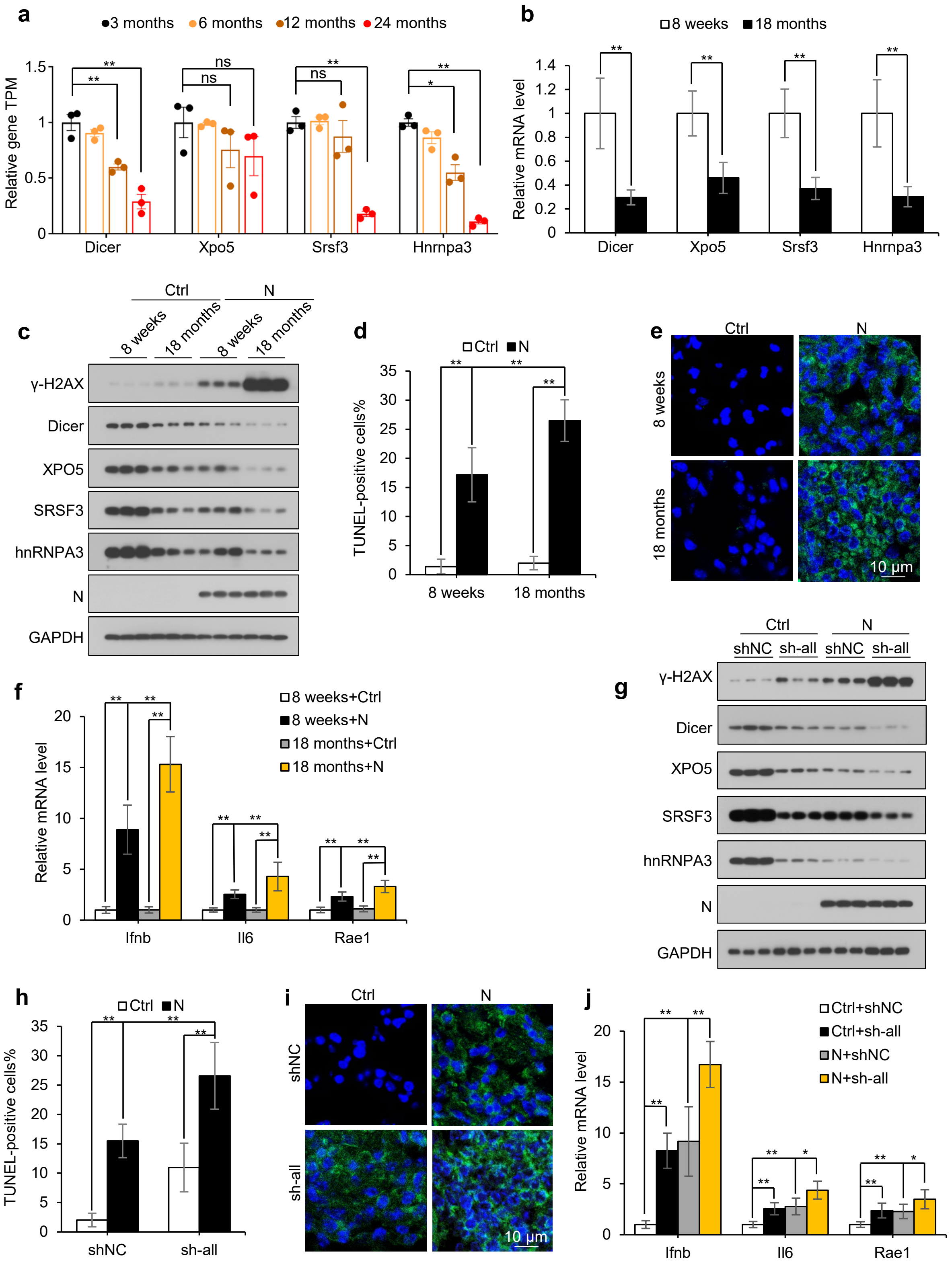

a
3 months 6 months 12 months 24 months
Relative gene TPM
Dicer Xpo5 Srsf3 Hnrnpa3
ns
b
8 weeks 18 months
Relative mRNA level
Dicer Xpo5 Srsf3 Hnrnpa3
c
Ctrl N
8 weeks 18 months 8 weeks 18 months
γ-H2AX
Dicer
XPO5
SRSF3
hnRNPA3
N
GAPDH
d
Ctrl N
TUNEL-positive cells%
8 weeks 18 months
e
Ctrl N
8 weeks
18 months
10 μm
f
8 weeks+Ctrl
8 weeks+N
18 months+Ctrl
18 months+N
Relative mRNA level
Ifnb Il6 Rae1
g
Ctrl N
shNC sh-all shNC sh-all
γ-H2AX
Dicer
XPO5
SRSF3
hnRNPA3
N
GAPDH
h
Ctrl N
TUNEL-positive cells%
shNC sh-all
i
Ctrl N
shNC
sh-all
10 μm
j
Ctrl+shNC
Ctrl+sh-all
N+shNC
N+sh-all
Relative mRNA level
Ifnb Il6 Rae1

**Extended Data Figure 6. Age-related downregulation of Dicer, XPO5, SRSF3, and hnRNPA3 expression is associated with N protein-induced pneumonia severity;** related to Fig. 5. **(a)** Expression levels of *Dicer*, *Xpo5*, *Srsf3*, and *Hnrnpa3* in the lung tissues of 3-month-, 6-month-, 12-month-, and 24-month-old mice. Data are downloaded from the Gene Expression Omnibus (GEO) database (GSE209891), n = 3 mice per group. $^{**}p < 0.01$; $^{*}p < 0.05$; ns, not significant ($p > 0.05$; two-sided Student's *t*-test). **(b)** mRNA levels of *Dicer*, *Xpo5*, *Srsf3*, and *Hnrnpa3* in the lung tissues of 8-week-old and 18-month-old mice. **(c–f)** Eight-week-old and 18-month-old mice were intranasally instilled with a control or N protein-expressing plasmid, and lung tissues were subjected to immunoblotting with the indicated antibodies (c), terminal deoxynucleotidyl transferase dUTP nick end labeling (TUNEL) assay (d), immunofluorescence with an anti-dsDNA antibody (e), and RT-qPCR analysis of *Ifnb*, *Il6*, and *Rae1* (f). **(g–j)** Eight-week-old mice intranasally instilled with N protein-expressing plasmid or co-instilled with N protein-expressing plasmid and plasmids expressing shRNAs targeting Dicer, XPO5, SRSF3, and hnRNPA3; lung tissues were subjected to immunoblotting with the indicated antibodies (g), TUNEL staining (h), immunofluorescence with anti-dsDNA antibody (i), and RT-qPCR analysis of *Ifnb*, *Il6*, and *Rae1* (j). Data in b, d, f, h, and j are expressed as mean ± SD of five mice. $^{**}p < 0.01$; $^{*}p < 0.05$; ns, not significant ($p > 0.05$; two-sided Student's *t*-test). Ctrl: control plasmid; N: N protein; shNC: negative control shRNA; sh-all: mice instilled with shDicer, shXPO5, shSRSF3, and shhnRNPA3 together.

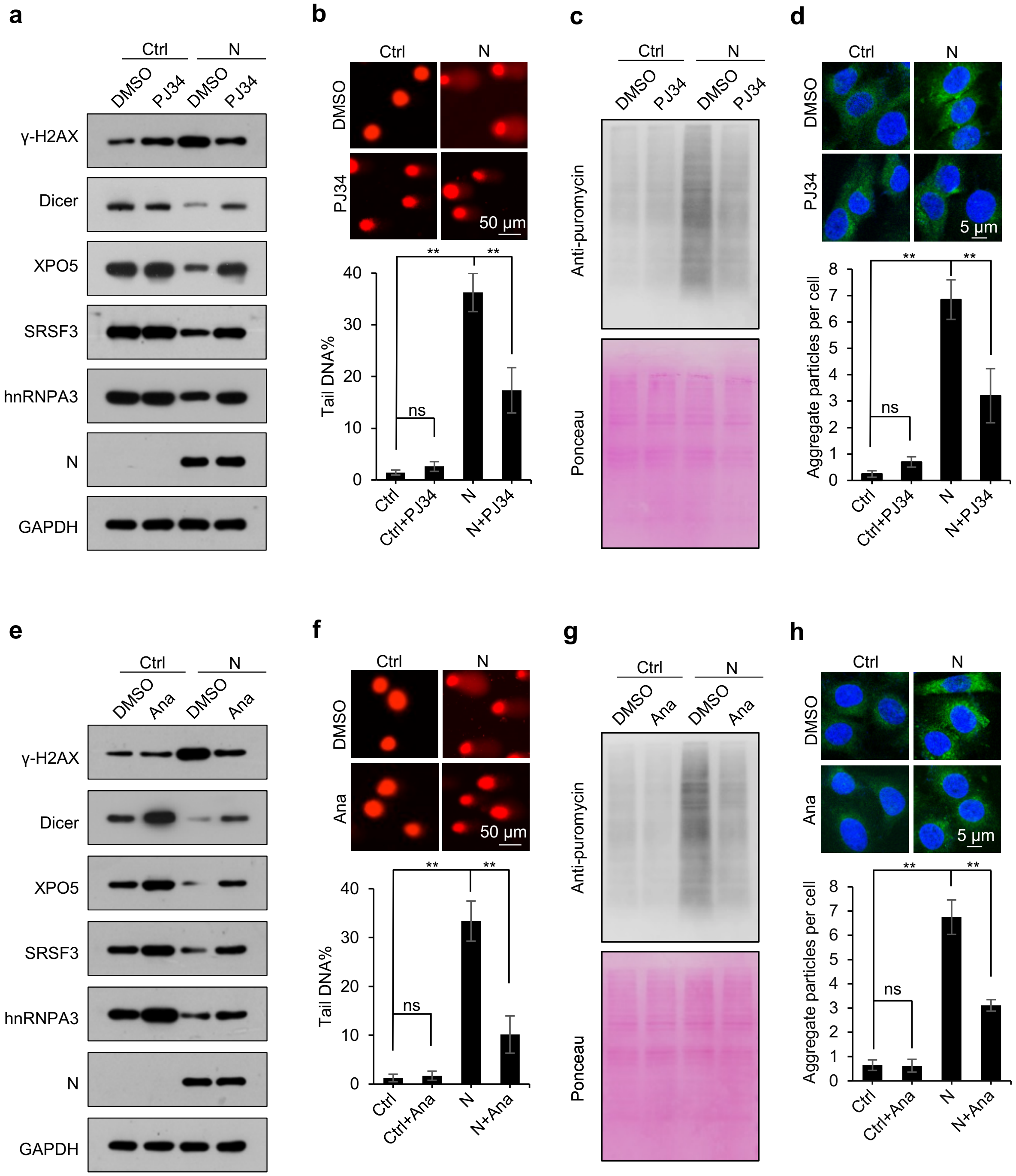
a
Ctrl
N
DMSO
PJ34
DMSO
PJ34
γ-H2AX
Dicer
XPO5
SRSF3
hnRNPA3
N
GAPDH
b
Ctrl
N
DMSO
PJ34
50 μm
Tail DNA%
ns
**
**
Ctrl
Ctrl+PJ34
N
N+PJ34
c
Ctrl
N
DMSO
PJ34
DMSO
PJ34
Anti-puromycin
Ponceau
d
Ctrl
N
DMSO
PJ34
5 μm
Aggregate particles per cell
ns
**
**
Ctrl
Ctrl+PJ34
N
N+PJ34
e
Ctrl
N
DMSO
Ana
DMSO
Ana
γ-H2AX
Dicer
XPO5
SRSF3
hnRNPA3
N
GAPDH
f
Ctrl
N
DMSO
Ana
50 μm
Tail DNA%
ns
**
**
Ctrl
Ctrl+Ana
N
N+Ana
g
Ctrl
N
DMSO
Ana
DMSO
Ana
Anti-puromycin
Ponceau
h
Ctrl
N
DMSO
Ana
5 μm
Aggregate particles per cell
ns
**
**
Ctrl
Ctrl+Ana
N
N+Ana

**Extended Data Figure 7. PJ34 and anastrozole relieve N protein-induced pneumonia;** related to Fig. 6. **(a)** Immunoblotting of the indicated proteins in BEAS-2B-Ctrl or BEAS-2B-N cells treated with or without PJ34 (50 μM) for 2 h. **(b)** Comet assay of DNA damage in A549-Ctrl or A549-N cells treated with or without PJ34 (50 μM) for 2 h. **(c)** Immunoblotting analysis of nascent polypeptides labeled with puromycin in A549-Ctrl or A549-N cells treated with or without PJ34 (50 μM) for 2 h. **(d)** Fluorescence image of A549 cells stably expressing the proteotoxic stress sensor reporters transfected with control or N protein-expressing plasmid in the presence or absence of PJ34 (50 μM). **(e)** Immunoblotting of the indicated proteins in BEAS-2B-Ctrl or BEAS-2B-N cells treated with or without anastrozole (5 μM) for 24 h. **(f)** DNA damage was assessed using comet assay in A549-Ctrl or A549-N cells treated with or without anastrozole (5 μM) for 24 h. **(g)** Immunoblotting analysis of nascent polypeptides labeled with puromycin in A549-Ctrl or A549-N cells treated with or without anastrozole (5 μM) for 24 h. **(h)** Fluorescence image of A549 cells stably expressing the proteotoxic stress sensor reporters transfected with control or N protein-expressing plasmid in the presence or absence of anastrozole (5 μM). Ponceau S staining images serve as the loading control in c and g. Data in b, d, f, and h are expressed as mean ± SD of three independent experiments. $^{**}p < 0.01$; ns, not significant ($p > 0.05$; two-sided Student's *t*-test). Ctrl: control plasmid; N: N protein; Ana: anastrozole.

#### Extended Data Tables

##### Extended Data Table 1. Plasmids used in this study.

| Name | Vendor | Catalog number |
|---|---|---|
| pLVX-EF1alpha-SARS-CoV-2-N-2xStrep-IRES-Puro | Addgene | 141391 |
| pLVX-EF1alpha-SARS-CoV-2-E-2xStrep-IRES-Puro | Addgene | 141385 |
| pLVX-EF1alpha-SARS-CoV-2-M-2xStrep-IRES-Puro | Addgene | 141386 |
| pLVX-EF1alpha-SARS-CoV-2-S-2xStrep-IRES-Puro | HITRO BioTech | Custom-made |
| pLVX-EF1alpha-2xStrep-IRES-Puro | HITRO BioTech | Custom-made |
| pLVX-3×FLAG-hACE2 | HITRO BioTech | Custom-made |
| CMV-LUC2CP/ARE | Addgene | 62857 |
| CMV-LUC2CP/intron/ARE | Addgene | 62858 |
| pCI-neo Fluc EGFP | Addgene | 90170 |
| pCI-neo FlucDM EGFP | Addgene | 90172 |
| pMD2.G | Addgene | 12259 |
| psPAX2 | Addgene | 12260 |
| pDESTmycDICER | Addgene | 19873 |
| pKmyc-Exp5 | Addgene | 12552 |
| pCDH-CMV-MCS-EF1-Puro-SRSF3(h) | Wuhan GeneCreate | Custom-made |
| pCDH-CMV-MCS-EF1-Puro-hnRNPA3(h) | Wuhan GeneCreate | Custom-made |
| pCDH-CMV-MCS-EF1-Puro-RNase H1(h) | Wuhan GeneCreate | Custom-made |
| pCDH-CMV-MCS-EF1-Puro-Dicer(m) | Wuhan GeneCreate | Custom-made |
| pCDH-CMV-MCS-EF1-Puro-XPO5(m) | Wuhan GeneCreate | Custom-made |
| pCDH-CMV-MCS-EF1-Puro-SRSF3(m) | Wuhan GeneCreate | Custom-made |
| pCDH-CMV-MCS-EF1-Puro-hnRNPA3(m) | Wuhan GeneCreate | Custom-made |
| pGL3-Control Vector | Promega | E1741 |
| pRL-CMV Vector | Promega | E2261 |
| pLKO.1-shNC-puro | Tsingke Biotech | Custom-made |
| pLKO.1-shFluc-puro | Tsingke Biotech | Custom-made |
| pLKO.1-shDicer(m)-puro | Tsingke Biotech | Custom-made |
| pLKO.1-shXPO5(m)-puro | Tsingke Biotech | Custom-made |
| pLKO.1-shSRSF3(m)-puro | Tsingke Biotech | Custom-made |
| pLKO.1-shhnRNPA3(m)-puro | Tsingke Biotech | Custom-made |

**Extended Data Table 2. Sequences (5′–3′) of siRNAs and primers used in this study.**

| Gene | siRNA and shRNA targeting sequences |
|---|---|
| shNC/siNC | TTCTCCGAACGTGTCACGT |
| shFluc | CTTACGCTGAGTACTTCGA |
| Human sip62-1 | GCATTGAAGTTGATATCGAT |
| Human sip62-2 | GGACCCATCTGTCTTCAAATT |
| Human siDicer-1 | AAGAGTTTACTAAGCACCAGG |
| Human siDicer-2 | AAGGCTTACCTTCTCCAGGCT |
| Human siXPO5-1 | GATGCTCTGTCTCGAATTGTA |
| Human siXPO5-2 | CCAGATGTTTCGAACACTAAA |
| Human siSRSF3-1 | AGAGCTAGATGGAAGAACATT |
| Human siSRSF3-2 | GCAACAAGACGGAATTGGATT |
| Human sihnRNPA3-1 | TCTTTACTTGTTAACTCACAA |
| Human sihnRNPA3-2 | GGAGGGAACTTTGGAGGTGTT |
| Mouse shDicer | AGATCACCGTCTCTAGAAA |
| Mouse shXPO5 | GATTTGATTTCGATAGTGATT |
| Mouse shSRSF3 | GGAAATAATGGAAACAAGAA |
| Mouse shhnRNPA3 | TTAAAGAGGATACGGAAGA |

| Gene | Forward/Reverse | Primer sequence |
|---|---|---|
| Human *IFNB* | Forward | TCTCCTCAGGGATGTCAAAG |
| | Reverse | CAACAAGTGTCTCCTCCAAAT |
| Mouse *Ifnb* | Forward | TCCTGCTGTGCTTCTCCACCACA |
| | Reverse | AAGTCCGCCCTGTAGGTGAGGTT |
| Human *IL6* | Forward | CAATCTGGATTCAATGAGGAGAC |
| | Reverse | CTCTGGCTTGTTCCTCACTACTC |
| Mouse *Il6* | Forward | GAGACTTCACAGAGGATACCAC |
| | Reverse | CAGTGCATCATCGCTGTTCATAC |
| Human *MICA* | Forward | CTTGGCCATGAACGTCAGG |
| | Reverse | CCTCTGAGGCCTCRCTGCG |
| Human *MICB* | Forward | ACCTTGGCTATGAACGTCACA |
| | Reverse | CCCTCTGAGACCTCGCTGCA |
| Human *ULBP1* | Forward | GTACTGGGAACAAATGCTGGAT |
| | Reverse | AACTCTCCTCATCTGCCAGCT |
| Human *ULBP2* | Forward | TTACTTCTCAATGGGAGACTGT |
| | Reverse | TGTGCCTGAGGACATGGCGA |
| Human *ULBP3* | Forward | CCTGATGCACAGGAAGAAGAG |

| | | |
|---|---|---|
| | Reverse | TATGGCTTTGGGTTGAGCTAAG |
| Human *ULBP4* | Forward | CGCCTTCTTTTGTTTCTGCTG |
| | Reverse | CCTGAGGTCTCGCCCCACT |
| Human *ULBP5* | Forward | CCTGGAAAGCACAGAACCCA |
| | Reverse | ACTGAGCTGCCAAGATCCAC |
| Human *ULBP6* | Forward | TCATCCCTAAGTTCAGACCTGG |
| | Reverse | GGACTGACGGGTGTGACTG |
| Mouse *H60* | Forward | GATGAACAGCATAGCATCTACT |
| | Reverse | CCTCATATCTTTCTCTAGGTTCT |
| Mouse *Rae1* | Forward | CAGTGACCAAGCGCCATCAT |
| | Reverse | ACCTAAGAGAGTGTGCATCATC |
| Human *GAPDH* | Forward | CTGGCGTCTTCACCACCATGG |
| | Reverse | CATCACGCCACAGTTTCCCGG |
| Mouse *Gapdh* | Forward | AACTTTGGCATTGTGGAAGG |
| | Reverse | CACATTGGGGGTAGGAACAC |
| Human *Dicer* | Forward | TCCACGAGTCACAATCAACACGG |
| | Reverse | GGGTTCTGCATTTAGGAGCTAGATGAG |
| Mouse *Dicer* | Forward | GCCAAGAAAATACCAGGTTGAGC |
| | Reverse | GCGATGAACGTCTTCCCTGAG |
| Human *XPO5* | Forward | ACGACGGGTGCATGGCTTCC |
| | Reverse | TTCGGCGCTTGTCAGCCACT |
| Mouse *Xpo5* | Forward | ACAAATTGCCATCGTCAGACA |
| | Reverse | CTCCAATCGGGACATGCTGT |
| Human *hnRNPA3* | Forward | GAAGGAGCTCTTCGCCTTTT |
| | Reverse | CAAACCTTACCCAGCCAGAA |
| Mouse *Hnrnpa3* | Forward | GAGGGCCATGATCCAAAGGAA |
| | Reverse | CACAAGAGTAGGTCACAAAACCA |
| Human *SRSF3* | Forward | AGCTGATGCAGTCCGAGAG |
| | Reverse | GGTGGGCCACGATTTCTAC |
| Mouse *Srsf3* | Forward | GACCACTCAGAAGTGTGTGGG |
| | Reverse | TCCTCAAATTCGACGAAAGCAAA |
| Human *ARGLU1* | Forward | AGCTGCTGATGCAGGTATTG |
| | Reverse | GTCCAGTGTCCTGCAGAGTG |
| Human *ARGLU1-IR* | Forward | GAAGAACTCGAGCGACAGAGA |
| | Reverse | CTGTTCTTCGGCCAGTTTG |
| Human *CCN1* | Forward | AAGATGCTTGTGGTTTGGCCC |
| | Reverse | TTCCGTATGCGCTTTCGTTG |

| | | |
|---|---|---|
| Human *CCN1-IR* | Forward | CAAGGGGCTGGAATGCAACT |
| | Reverse | TTCTCGTCAACTCCACCTCG |
| Human *TERT* | Forward | ACCTTCCTCAGGACCCTGGT |
| | Reverse | ATCTGAACAAAAGCCGTGCC |
| Human *TERT-IR* | Forward | GCCAATCCCAAAGGGTCAGA |
| | Reverse | TCGGGTTCAGAGGGACTCAT |

**Extended Data Table 3. Antibodies used in this study.**

| Antibody | Vendor | Catalog number |
|---|---|---|
| Rat anti-Strep-tag II | Abcam | ab252885 |
| Rabbit anti-SARS-CoV-2 (2019-nCoV) Nucleocapsid | Sino Biological Inc. | 40588-RC02 |
| Rabbit anti-ACE2 | Proteintech | 21115-1-AP; RRID:AB_10732845 |
| Rabbit anti-SRSF3 | Abcam | ab198291 |
| Mouse anti-Dicer | Abcam | ab14601; RRID:AB_443067 |
| Rabbit anti-Exportin 5 | Cell Signaling Technology | 12565; RRID:AB_2737081 |
| Rabbit anti-RNase H1 | Proteintech | 15606-1-AP; RRID:AB_2238624 |
| Rabbit anti-hnRNPA3 | Proteintech | 25142-1-AP; RRID:AB_2879921 |
| Rabbit anti-Phospho-ATM (Ser1981) | Cell Signaling Technology | 13050; RRID:AB_2798100 |
| Rabbit anti-ATM | Proteintech | 27156-1-AP; RRID:AB_2880780 |
| Rabbit anti-Phospho-ATR (Ser428) | Cell Signaling Technology | 2853; RRID:AB 2290281 |
| Rabbit anti-ATR | Proteintech | 19787-1-AP; RRID:AB_10639516 |
| Rabbit anti-Phospho-Chk1 (Ser345) | Cell Signaling Technology | 2348; RRID:AB_331212 |
| Rabbit anti-Chk1 | Proteintech | 25887-1-AP; RRID:AB_2880283 |
| Rabbit anti-Phospho-Chk2 (Thr68) | Cell Signaling Technology | 2661; RRID:AB_331479 |
| Rabbit anti-Chk2 | Cell Signaling Technology | 2662; RRID:AB_2080793 |
| Rabbit anti-Phospho-Histone H2A.X (Ser139) | Cell Signaling Technology | 2577; RRID:AB_2118010 |
| Rabbit anti-Histone H2A.X Polyclonal | Proteintech | 10856-1-AP; RRID:AB_2114985 |
| Mouse anti-SQSTM1/p62 | Cell Signaling Technology | 88588; RRID:AB_2800125 |
| Rabbit anti-SQSTM1/p62 | Proteintech | 18420-1-AP; RRID:AB_10694431 |
| Rabbit anti-LC3 | Proteintech | 14600-1-AP; RRID:AB_2137737 |
| Rabbit anti-Phospho-SAPK/JNK | Cell Signaling Technology | 9251; |

| | | |
|---|---|---|
| (Thr183/Tyr185) | | RRID:AB_331659 |
| Rabbit anti-JNK1/2/3 | Proteintech | 28007-1-AP; RRID:AB_2881035 |
| Rabbit anti-Phospho-p70(S6K) (Thr389) | Proteintech | 28735-1-AP; RRID:AB_2918197 |
| Mouse anti-p70(S6K) | Proteintech | 66638-1-Ig; RRID:AB_2881997 |
| Rabbit anti-Phospho-4EBP1 (Ser65/Thr70) | Abmart | TA2308 |
| Mouse anti-4EBP1 | Proteintech | 60246-1-Ig; RRID:AB_2881368 |
| Rabbit anti-CD3 | Proteintech | 17617-1-AP; RRID:AB_1939430 |
| Rabbit anti-CD22 | Proteintech | 21894-1-AP; RRID:AB_2878937 |
| Rabbit anti-CD68 | Proteintech | 28058-1-AP; RRID:AB_2881049 |
| Rabbit anti-Ly6G | Abcam | ab238132; RRID: AB_2923218 |
| Mouse anti-GAPDH | Proteintech | 60004-1-Ig; RRID:AB_2107436 |
| Mouse anti-DNA antibody, double stranded, clone AE-2 | Sigma-Aldrich | MAB1293; RRID:AB_11215105 |
| Mouse anti-puromycin | Sigma-Aldrich | MABE343; RRID:AB_2566826 |
| Mouse anti-DNA-RNA hybrid (R-loop) antibody, clone S9.6 | Sigma-Aldrich | MABE1095; RRID:AB_2861387 |
| Rabbit IgG | Beyotime Biotechnology | A7058 |
| Mouse IgG | Beyotime Biotechnology | A7050 |
| Mouse anti-rabbit IgG (Conformation Specific) (L27A9) mAb (HRP Conjugate) | Cell Signaling Technology | 5127; RRID:AB_10892860 |
| Goat anti-rabbit IgG (H+L) secondary antibody, HRP conjugate | Biosharp | BL003A; RRID:AB_2827666 |
| Goat anti-mouse IgG (H+L) secondary antibody, HRP conjugate | Biosharp | BL001A; RRID:AB_2827665 |
| Goat anti-Rabbit IgG (H+L) Cross-Adsorbed Secondary Antibody, Alexa Fluor™ 488 | Thermo Fisher Scientific | A-11008; RRID:AB_143165 |
| Rabbit anti-Rat IgG (H+L) secondary antibody, HRP conjugate | Boster | BA1058; RRID:AB_10891896 |
| Goat anti-Rabbit IgG (H+L) Cross-Adsorbed Secondary Antibody, Alexa Fluor™ 555 | Thermo Fisher Scientific | A-21428; RRID:AB_2535849 |
| Goat anti-Mouse IgG (H+L) Cross-Adsorbed Secondary Antibody, Alexa Fluor™ 488 | Thermo Fisher Scientific | A-11001; RRID:AB_2534069 |
